# Regulation of Mus81-Eme1 structure-specific endonuclease by Eme1 SUMO-binding and Rad3^ATR^ kinase is essential in the absence of Rqh1^BLM^ helicase

**DOI:** 10.1101/2021.07.29.454171

**Authors:** C. Giaccherini, S. Scaglione, S. Coulon, P.M. Dehé, P.H.L. Gaillard

## Abstract

The Mus81-Eme1 structure-specific endonuclease is crucial for the processing of DNA recombination and late replication intermediates. In fission yeast, stimulation of Mus81-Eme1 in response to DNA damage at the G2/M transition relies on Cdc2^CDK1^ and DNA damage checkpoint-dependent phosphorylation of Eme1 and is critical for chromosome stability in absence of the Rqh1^BLM^ helicase. Here we identify Rad3^ATR^ checkpoint kinase consensus phosphorylation sites and two SUMO interacting motifs (SIM) within a short N-terminal domain of Eme1 that is required for cell survival in absence of Rqh1^BLM^. We show that catalytic stimulation of Mus81-Eme1 depends entirely on direct phosphorylation of Eme1 by Rad3^ATR^ and that while Eme1 also undergoes Chk1-mediated phosphorylation, this is not essential for catalytic modulation. Both Rad3^ATR^- and Chk1-mediated phosphorylation of Eme1 as well as the SIMs are independently critical for cell fitness in absence of Rqh1^BLM^ and abrogating bimodal phosphorylation of Eme1 along with mutating the SIMs is incompatible with *rqh1Δ* cell viability. Our findings unravel an elaborate regulatory network that relies on the poorly structured N-terminal domain of Eme1 and which is essential for the vital functions Mus81-Eme1 fulfills in absence of Rqh1^BLM^.

## Introduction

Structure-specific DNA endonucleases are cornerstones in the proper execution of DNA replication, repair and recombination; yet they harbor the potential for causing genome instability. Controlling these enzymes is essential to ensure efficient processing of appropriate substrates while preventing counterproductive targeting of other similar DNA structures. The Mus81-Eme1 structure-specific endonuclease (SSE) has emerged as a key player in the processing of recombination intermediates that form during homology directed repair of two-ended double strand breaks or during the rescue of stalled or broken replication forks. Over this last decade, elaborate control mechanisms of Mus81-Eme1 that are tightly linked to cell cycle progression have been identified.

In *Saccharomyces. cerevisiae* (*S. cerevisiae*), Mus81-Mms4^EME1^ is stimulated at the G2/M transition by Cdc5^PLK1^-, Cdc28^CDK1^- and Dbf4-dependent phosphorylation of Mms4^EME1^ [1–5]. In human cells, catalytic upregulation of MUS81-EME1 is driven by complex formation with the SLX4 nuclease scaffold[6–8]. This regulation is mediated by direct interaction between MUS81 and SLX4, which is strongly stimulated at the G2/M transition by phosphorylation of SLX4 by CDK1[9]. Reminiscent of what has been observed in *S. cerevisiae*, maximal processing of joint molecules such as Holliday junctions by MUS81-EME1 also correlates with hyperphosphorylation of EME1, possibly by CDK1 or PLK1[4]. Whether phosphorylation of EME1 contributes to the stimulation of MUS81-EME1 in human cells remains to be formally demonstrated. These mechanisms ensure that joint molecules such as Holliday junctions, D-loops or replication intermediates at under-replicated loci are efficiently resolved in mitosis before chromosome segregation[10]. By restricting the catalytic stimulation of Mus81-Eme1 to late stages of the cell cycle these control mechanisms also ensure that joint molecules and replication intermediates get a chance to be processed by more conservative non-endonucleolytic mechanisms that rely on their unfolding by RecQ helicases such as the BLM helicase in human cells and its Sgs1 and Rqh1^BLM^ orthologs in *S. cerevisiae* and *Schizosaccharomyces pombe* (*S. pombe*), respectively. These temporal controls further prevent the accumulation of hyper-activated Mus81-Eme1 in S-phase and the risk of the unscheduled processing of replication intermediates. The importance of such control mechanisms is underscored by the marked genomic instability that is caused by the premature stimulation of Mus81-Mms4 in budding yeast cells that produce an Mms4 mutant that mimics a constitutively phosphorylated Mms4 protein[2]. Interestingly, SUMOylation and ubiquitination of Mms4 were recently shown to specifically target phosphorylated Mms4 for degradation by the proteasome, further ensuring that hyperactivation of Mus81-Mms4 is restricted to mitosis[11]. In human cells, SLX4-MUS81 complex formation induced by premature activation of CDK1 results in the unscheduled processing of replication intermediates genome wide and chromosome pulverization[9].

In *S. pombe*, upregulation of Mus81-Eme1 also relies on the phosphorylation of Eme1 by Cdc2^CDK1^[12]. However, in contrast to what has been described in *S. cerevisiae*, phosphorylation of Eme1 by Cdc2^CDK1^ primes Eme1 for further DNA damage checkpoint-mediated phosphorylation in response to DNA damage. This elaborate control mechanism ensures that Mus81-Eme1 is rapidly hyperactivated in response to DNA damage in late G2 and mitosis and is critical to prevent gross chromosomal rearrangements in absence of the BLM-related helicase Rqh1^BLM^[12].

To gain further insight into the molecular mechanisms involved in the control of Mus81-Eme1 in *S. pombe*, we undertook *in silico* analyses of a relatively short N-terminal domain of Eme1 that we found to be essential for cell viability in the absence of Rqh1^BLM^. We identified Rad3^ATR^ consensus phosphorylation sites and two SUMO interacting Motifs (SIM1 and SIM2) within that domain. We demonstrate that Eme1 is a direct substrate for Rad3^ATR^ both *in vitro* and *in vivo* and show that it is phosphorylation of Eme1 by Rad3^ATR^, not Chk1 as initially proposed, that primarily leads to the catalytic stimulation of Mus81-Eme1 in response to DNA damage. We provide genetic evidence that both Rad3^ATR^ - and Chk1-mediated phosphorylation of Eme1 are independently critical for cell fitness in the absence of Rqh1^BLM^. Remarkably, Chk1-mediated phosphorylation of Eme1 is lost when SIM2 is mutated, while mutating SIM1 has no impact. Both SIMs are important for cell fitness in absence of Rqh1^BLM^, while abrogating phosphoregulation of Mus81-Eme1 and mutating SIM1 and SIM2 recapitulates the synthetic lethality observed by deleting the N-terminus of Eme1 in the absence of Rqh1^BLM^.

## Results

### Eme1 N-terminus is essential in absence of Rqh1^BLM^

To gain further insight into the mechanisms underlying the catalytic stimulation of Mus81-Eme1 in response to DNA damage, we searched for domains of Eme1 that are essential in absence of Rqh1^BLM^ while dispensable for the intrinsic catalytic activity of Mus81-Eme1. Interestingly, we found that deleting a relatively short N-terminal domain (residues 1-117) of Eme1 is synthetic lethal with *rqh1Δ* (**Fig 1A**). Importantly, this domain does not contain the Cdc2^CDK^ sites that we had previously reported to be involved in the stimulation of Mus81-Eme1 in response to DNA damage and to be critical for cell viability in absence of Rqh1^BLM^ [12]. A detailed *in silico* analysis of the first 117 residues of Eme1 led to the identification of two SQ/TQ Rad3^ATR^ consensus sites (S23Q and T50Q) and two putative SUMO-Interacting Motifs (SIMs), hereafter named SIM1 and SIM2, which matched the described (V/I)-X-(V/I)-X-(V/I/L) consensus sequence (**Fig 1B**)[13]. These observations suggested that Eme1 might be a direct substrate for Rad3^ATR^ and possess SUMO-binding properties. We tested these predictions in the following experiments.

**Figure 1:**
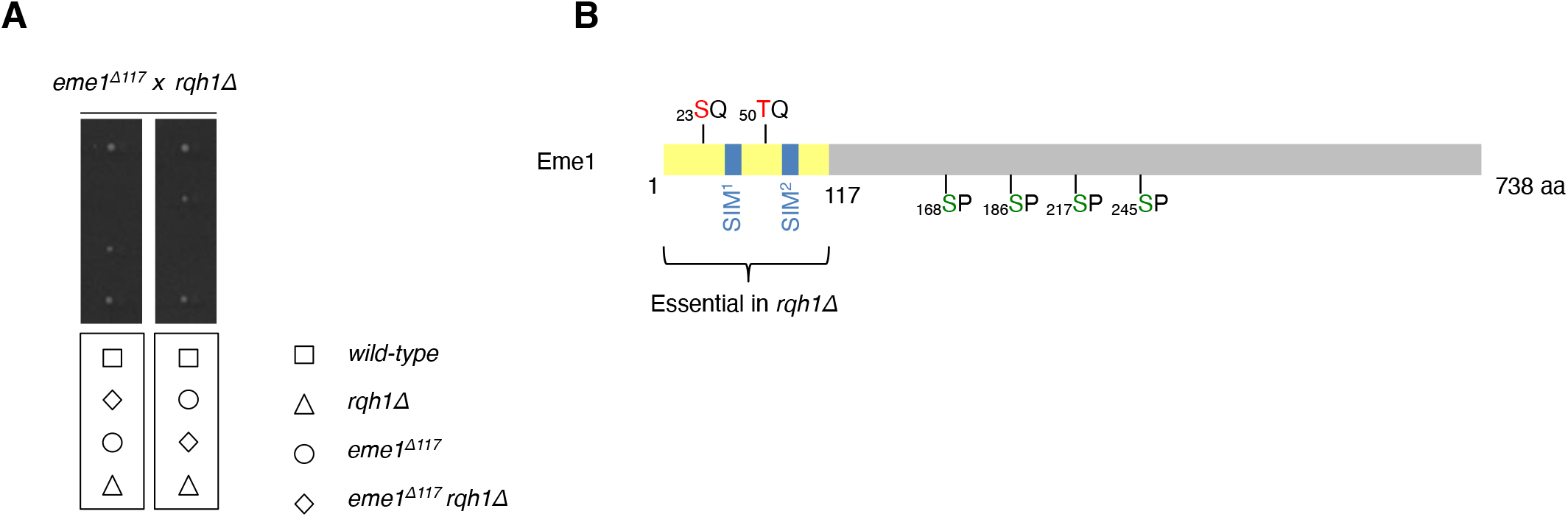
Eme1 N-terminal part (1-117) is essential in absence of Rqh1^BLM^. A- Tetrad analysis of an *eme1^Δ117^* x *rqh1Δ* mating, germinated at 30 °C. Boxes below dissections indicate the genotypes of each spore. B- Schematic of Eme1 protein. The yellow box depicts the first 117 amino-acid residues required for cell survival in absence of Rqh1^BLM^. The serine and threonine residues responding to Rad3^ATR^-consensus phosphorylation sites are depicted in red and the SUMO-Interacting Motifs are represented by blue squares. The serine residues targeted by Cdc2 are depicted in green.

### Eme1 is a direct target of Rad3^ATR^ kinase

Eme1 contains in total eight putative Rad3^ATR^ consensus phosphorylation sites (S_23_Q, T_50_Q, S_126_Q, T_145_Q, T_215_Q, S_313_Q, T_384_Q, S_458_Q) (**Fig S1**). To determine whether Eme1 is a direct substrate of Rad3^ATR^ we set up *in vitro* kinase assays with recombinant Mus81-Eme1 and Rad3^ATR^. Recombinant Mus81(6His)-(MBP)Eme1 was produced in *E. coli* and affinity purified on Ni++ and amylose resins (**Fig 2A**). Recombinant (GFP)Rad3^ATR^ was instead transiently overproduced in yeast cells exposed to bleomycin in order to induce DNA damage and activate Rad3^ATR^. We used a *chk1Δ cds1Δ rad3Δ* mutant strain to eliminate the possibility of endogenous checkpoint kinases co-purifying with (GFP)Rad3^ATR^. As shown in **Figure 2B,** Eme1 was efficiently phosphorylated by (GFP)Rad3^ATR^. To explore which SQ/TQ sites were important for phosphorylation we produced Mus81-Eme1 complexes in which SQ/TQ sites were mutated to AQ in three different clusters (**Fig S1**). The strongest effect was seen with mutations in Cluster 1, whilst mutations in Clusters 2 and 3 had a mild impact (**Fig 2B and C**). Noticeably, *in vitro* Rad3^ATR^-mediated phosphorylation of Eme1 does not require the priming by Cdc2^CDK1^. Taken together, these data indicate that the S_23_Q and T_50_Q sites in the N-terminus of Eme1 are most critical for its *in vitro* phosphorylation by Rad3^ATR^.

**Figure 2:**
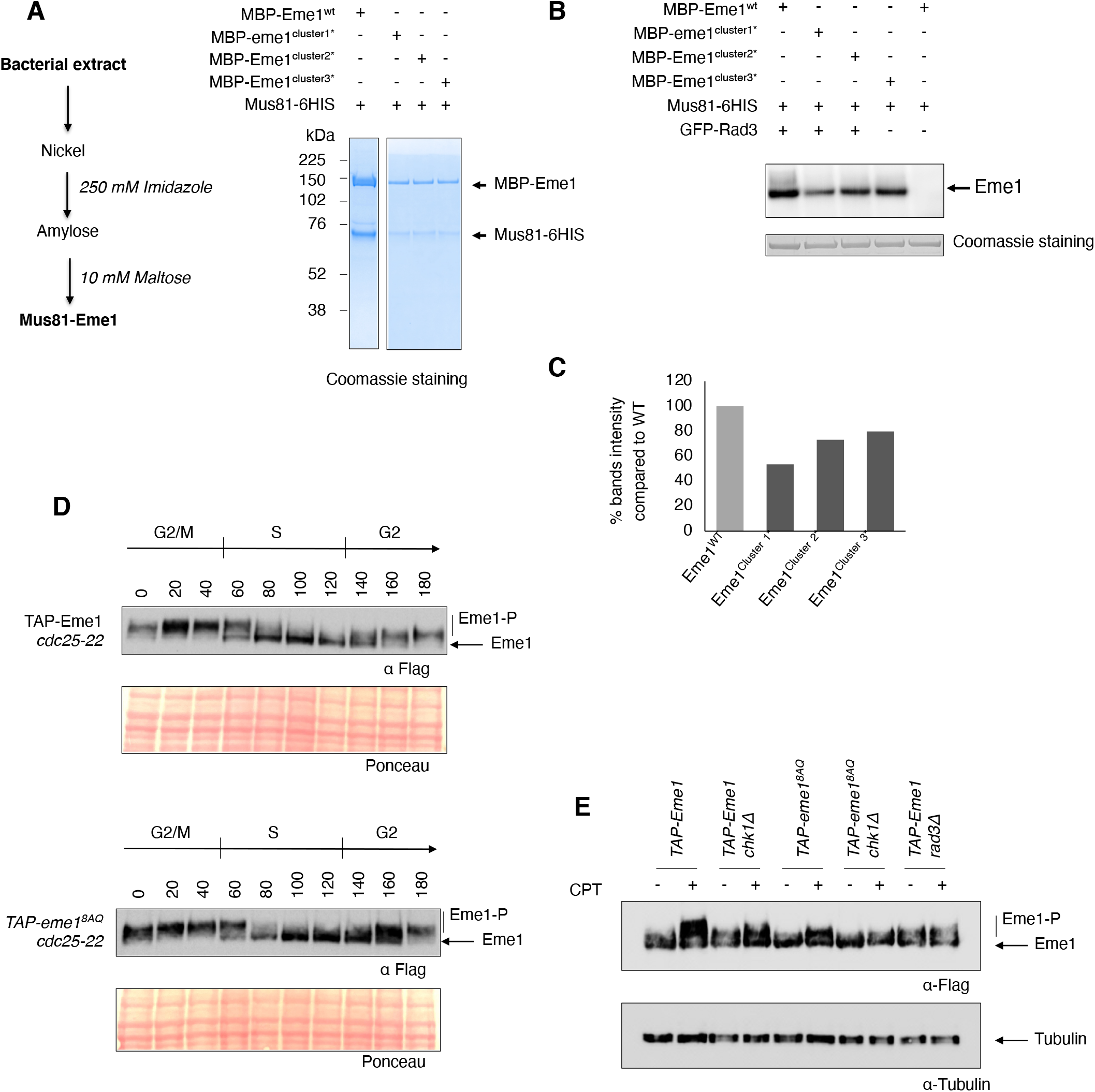
Rad3^ATR^ directly phosphorylates Eme1 *in vitro*. A- Recombinant *mus81* was co-expressed with either wild-type *eme1, eme1^cluster1*^, eme1^cluster2*^* or *eme1^cluster3*^* and purified from bacterial cultures (purification scheme is depicted on the left). B- Rad3^ATR^ in vitro kinase assay on full-length recombinant Mu81-Eme1 complexes containing either wild-type Eme1 or Eme1 mutated for S_23_/T_50_ (cluster1*) or S_126_/T_145_/T_215_ (cluster 2*) or S_313_/T_384_/S_458_ (cluster 3*). C- All clusters contribute to Eme1 Rad3^ATR^-dependent phosphorylation with S_23_/T_50_ (cluster1*) being the most important. D- Cultures from *cdc25-22 TAP-eme1* and *cdc25-22 TAP-eme1^8AQ^* were synchronized at the G2/M transition and released for one cell cycle. Total proteins were extracted at each indicated time point of the time course and analyzed by Western blot using an antibody raised against the Flag tag of Eme1. Ponceau stained membranes are depicted as loading control. E- Western blot on total lysates from untreated or 40 μM CPT-treated cells of the indicated background. Tubulin is used as a loading control. Note: TAP- = 2xProtA-TEVsite-2xFlag-

To investigate whether Eme1 is a substrate for Rad3^ATR^ *in vivo*, we generated an *eme1^8AQ^* mutant strain in which all SQ/TQ sites are mutated to AQ (**Fig S1**). As expected, whereas the cell-cycle dependent phosphorylation profile of Eme1^8AQ^ is comparable to that of the WT protein (**Fig 2D**), we observed a strong reduction of the CPT-induced phosphorylation (**Fig 2E**). However, Rad3^ATR^-dependent phosphorylation was not totally abolished in the Eme1^8AQ^ background (**Fig 2E**). We suspected that the residual Rad3^ATR^-dependent phosphorylation of Eme1^8AQ^ was catalyzed by Chk1. Accordingly, we observed a complete loss of CPT-induced phosphorylation of Eme1^8AQ^ in *eme1^8AQ^ chk1Δ* cells (**Fig 2E**).

Overall, our data strongly indicate that Eme1 is phosphorylated by both Chk1 and Rad3^ATR^ following activation of the DNA damage checkpoint.

### Phosphorylation of Eme1 by Rad3^ATR^ is crucial in absence of Rqh1^BLM^

We previously showed that *in vivo* DNA damage-induced hyperphosphorylation of Eme1 is strictly subordinated to prior Cdc2^CDK1^-dependent phosphorylation [12]. Accordingly, mutating four CDK consensus target sites totally abrogates not only the cell-cycle dependent phosphorylation of the resulting Eme1^4SA^ protein but also its phosphorylation in response to DNA damage[12]. Importantly, the same study found that while an *eme1^4SA^* single mutant displays no abnormal phenotype, an *eme1^4SA^ rqh1Δ* double mutant is extremely sick. These observations suggest that the Cdc2^CDK1^-dependent phosphorylations of Eme1 are required for its phosphorylation by Rad3^ATR^, and these events are critical in the absence of Rqh1^BLM^.

To investigate the functional relevance of phosphorylation of Eme1 by Rad3^ATR^, we introduced *eme1^8AQ^* mutations in the *rqh1Δ* background. While we observed no obvious phenotype in the *eme1^8AQ^* single mutant, the *eme1^8AQ^ rqh1Δ* double mutant displayed pronounced growth and colony formation defects compared to the *rqh1Δ* single mutant (**Fig 3A-C and S2A**). This genetic interaction was further exacerbated by exposure to genotoxic agents (**Fig S2B**). Hence, direct phosphorylation of Eme1 by Rad3^ATR^ appears to be important for cell viability in the absence of Rqh1^BLM^.

**Figure 3:**
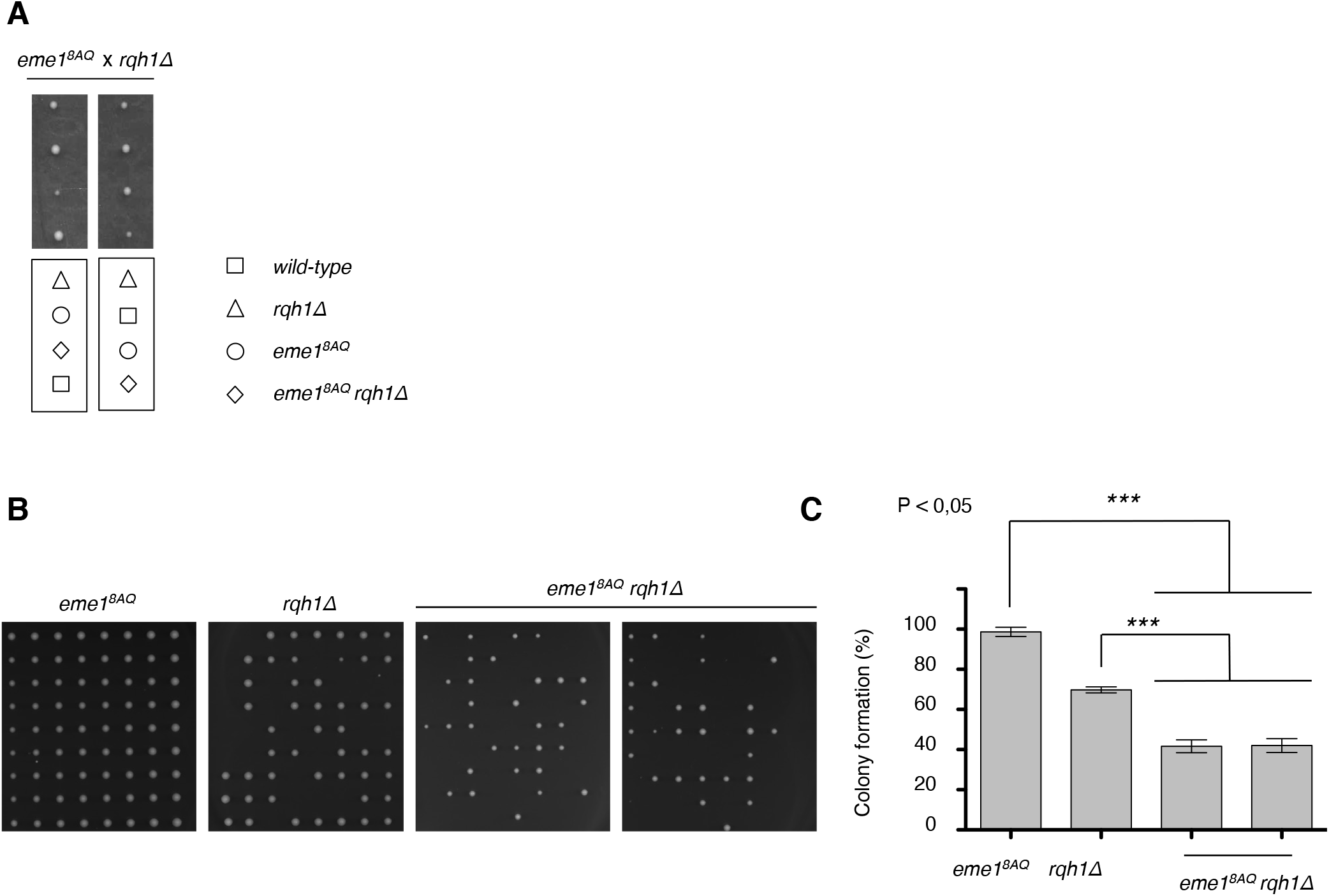
Negative genetic interaction between *eme1^8AQ^* and *rqh1Δ*. A- Tetrad analysis of an *eme1^8AQ^* x *rqh1Δ* mating, germinated at 30 °C. Boxes below dissections indicate the genotypes of each spore. B- Exponentially growing *eme1^8AQ^, rqh1Δ* and *eme1^8AQ^ rqh1Δ* cells were seeded on YES plates by micromanipulation and allowed to grow for 3 days at 32°C. C- Average percentage (± s.d.) of viable colonies (n=3 independent experiments). Statistical significance is measured with one-way ANOVA followed by Tuckey post-test.

### Rad3^ATR^ direct phosphorylation of Eme1 contributes to the catalytic stimulation of Mus81-Eme1

We have previously shown that Rad3^ATR^ contributes to the catalytic stimulation of the HJ-resolvase activity of Mus81-Eme1[12]. We inferred at that time that the Rad3^ATR^-Chk1 axis was involved in this catalytic control. The finding that Rad3^ATR^ can directly phosphorylate Eme1, and this phosphorylation is crucial in absence of Rqh1^BLM^, prompted us to investigate whether it contributed to the catalytic stimulation of Mus81-Eme1.

As previously reported[12], hyperphosphorylation of Eme1 following activation of the DNA damage checkpoint by CPT correlates with increased HJ-resolvase activity of Mus81-Eme1 complex isolated from fission yeast cells expressing TAP-tagged Eme1 (**Fig 4A and S3A**). Accordingly, no catalytic stimulation is detected when Mus81-Eme1 is precipitated from *rad3Δ* cells ([12] and **Fig 4B and S3B**). Remarkably, we find that the Mus81-Eme1^8AQ^ complex is not stimulated following CPT treatment despite a fully functional DNA damage checkpoint in *eme1^8AQ^* mutant (**Fig 4C and S3C**). This demonstrates that it is the phosphorylation of Eme1 by Rad3^ATR^, rather than by Chk1, that contributes to the catalytic stimulation of Mus81-Eme1. Consistently, Mus81-Eme1 can still be stimulated in *chk1Δ* cells following CPT treatment (**Fig 4D and S3D**). Overall, our data demonstrate that catalytic stimulation of Mus81-Eme1 relies primarily on phosphorylation of Eme1 by Rad3^ATR^ and not Chk1.

**Figure 4:**
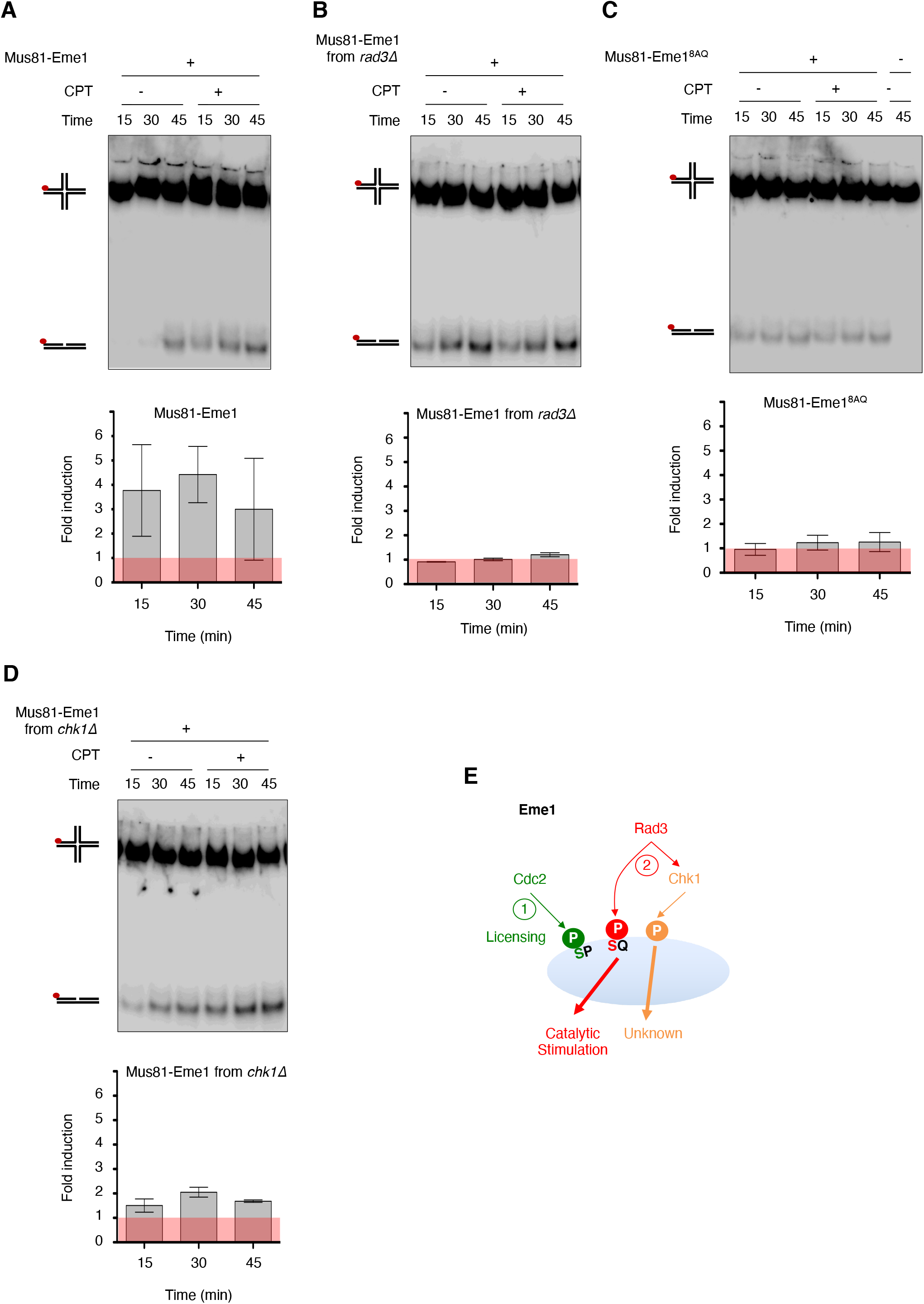
Rad3^ATR^ phosphorylation of Eme1 contributes to the catalytic stimulation of Mus81-Eme1 resolvase activity. A- Upper panel: Neutral PAGE showing ^32^P-labeled (red dot) HJs incubated for the indicated times with Mus81-Eme1 complexes recovered from untreated or 40 μM CPT-treated *TAP-eme1* (“wild type”) cells as described in Materials and Methods. Identical amounts of TEV eluates were used in each reaction after normalization of their relative concentration (see Materials and Methods). Lower panel: Average (± s.d.) fold stimulation of HJ resolution by Mus81-Eme1 following CPT-treatment (n=3 independent experiments, see S3). The histogram shows the ratio of the HJ-resolvase activity of Mus81-Eme1 from CPT-treated *TAP-eme1* (“wild type”) cells over that of Mus81-Eme1 from untreated *TAP-eme1* (“wild type”) cells (fold induction). B- Upper panel: Same as Upper panel A- but with Mus81-Eme1 from *TAP-eme1 rad3Δ* cells. Lower panel: Same as Lower panel A- but with Mus81-Eme1 from *TAP-eme1 rad3Δ* cells. C- Upper panel: Same as Upper Panel A- but with Mus81-Eme1 from *TAP-eme1 eme1^8AQ^* cells. Lower panel: Same as Lower panel A- but with Mus81-Eme1 from *TAP-eme1 eme1^8AQ^* cells. D- Upper panel: Same as Upper panel A- but with Mus81-Eme1 from *TAP-eme1 chk1Δ* cells. Lower panel: Same as Lower panel A- but with Mus81-Eme1 from *TAP-eme1 chk1Δ* cells. E- Model for the Cdc2-, Rad3^ATR^- and Rad3^ATR^-Chk1-dependent phosphorylation of Eme1 and their respective contributions. Note: TAP- = 2xProtA-TEVsite-2xFlag-

Here we have shown that two axes phosphorylate Eme1 in response to DNA damage with different outcomes. The Rad3^ATR^-Chk1 branch of the DNA damage checkpoint phosphorylates Eme1 and contributes to non-catalytic functions of the complex while Rad3^ATR^ direct phosphorylation of Eme1 stimulates the HJ-resolvase activity of the Mus81-Eme1 complex (**Fig 4E**).

### Eme1 contains bona-fide SIMs

Having confirmed our predictions that Eme1 is directly phosphorylated by Rad3^ATR^ and shown that this is critical for catalytic stimulation of Mus81-Eme1 and cell fitness in absence of Rqh1^BLM^, we next undertook the analysis of the predicted SUMO-binding properties of Eme1 mediated by the putative SIM1 and SIM2 motifs in the N-terminal domain of Eme1 (**Fig 1B and 5A**).

**Figure 5:**
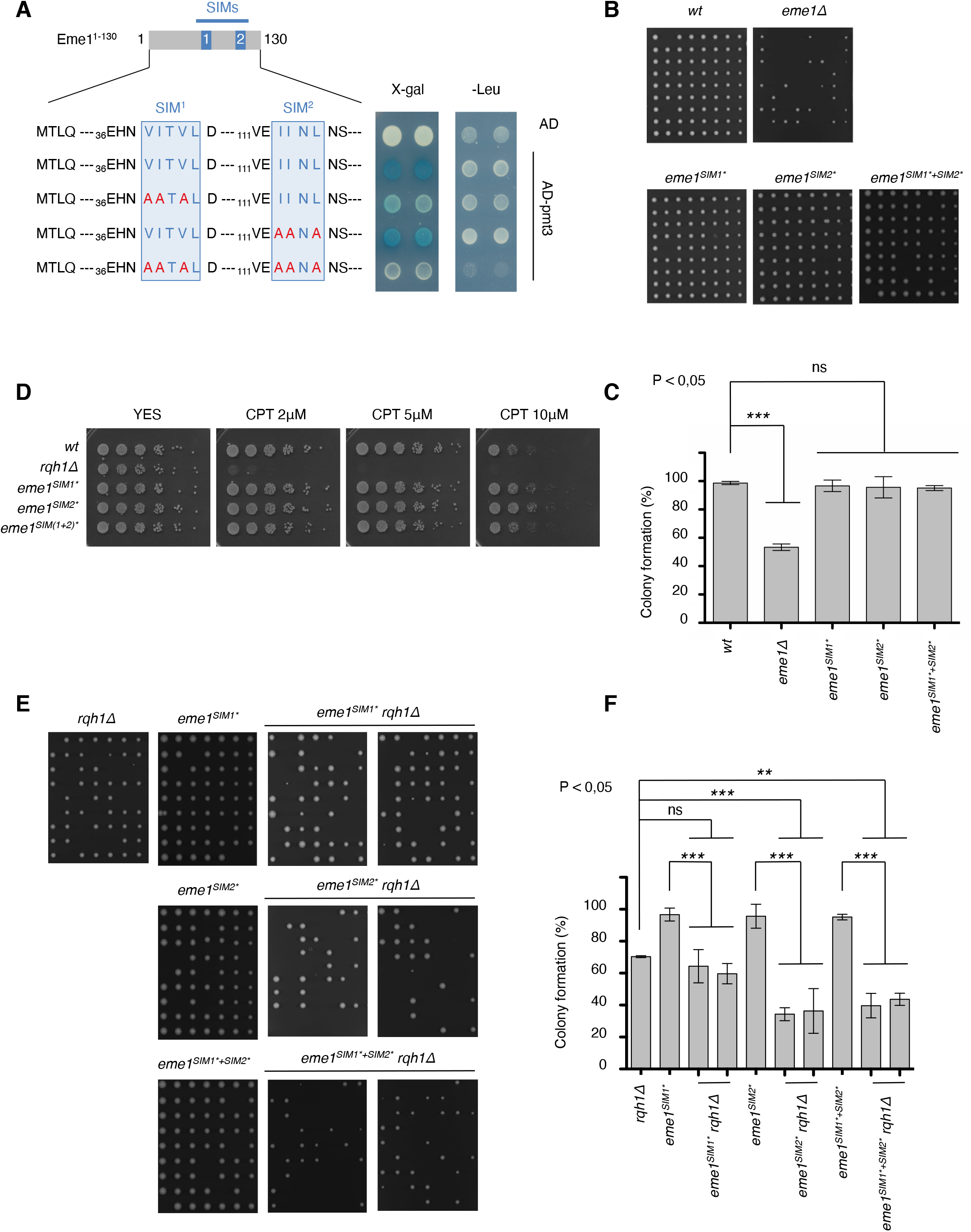
Eme1 contains *bona-fide* SIMs. A- A fragment of Eme1 (eme1^1-130^) containing SIM1 and SIM2 was used as bait in a Yeast two-hybrid assay for interaction with SUMO^PMT3^. The SIMs consensus sequences are indicated as well as the mutations introduced in each of them. B- Exponentially growing wild type, *eme1Δ, eme1^SIM1*^, eme1^SIM2*^* and *eme1^SIM1*+SIM2*^* cells were seeded on YES plates by micromanipulation and allowed to grow for 3 days at 32°C. C- Average percentage (± s.d.) of viable colonies (n=3 independent experiments). Statistical significance is measured with one-way ANOVA followed by Tuckey post-test. D- Five-fold dilutions of wild-type, *rqh1Δ, eme1^SIM1*^, eme1^SIM2*^* and *eme1^SIM1*+SIM2*^* cells were plated on medium supplemented or not with the indicated concentrations of CPT followed by incubation at 30 °C. E- Exponentially growing *rqh1Δ, eme1^SIM1*^, eme1^SIM2*^* and *eme1^SIM1*+SIM2*^* as well as two independent clones of *eme1^SIM1*^ rqh1Δ, eme1^SIM2*^ rqh1Δ* and *eme1^SIM1*+SIM2*^ rqh1Δ* were seeded on YES plates by micromanipulation and allowed to grow for 3 days at 32°C. F- Average percentage (± s.d.) of viable colonies (n=3 independent experiments). Statistical significance is measured with one-way ANOVA followed by Tuckey post-test.

The SUMO-binding capacities of SIM1 and SIM2 were assessed by a yeast two-hybrid assay against the unique *S. pombe* SUMO ortholog Pmt3. A fragment of Eme1 (Eme1^1-130^) containing SIM1 and SIM2 displayed strong binding to Pmt3, confirming that the N-terminus of Eme1 possesses SUMO-binding properties (**Fig 5A**). Introducing point mutations in the conserved aliphatic residues of SIM1 strongly impaired interaction with Pmt3 (**Fig 5A**) while mutations in SIM2 had a milder effect (**Fig 5A**). Mutations in both SIMs led to complete loss of interaction with Pmt3 (**Fig 5A**). Our data confirm that the N-terminal domain of Eme1, which is essential for cell viability in absence of Rqh1^BLM^ (**Fig 1A**), contains *bona fide* SIMs that jointly contribute to the SUMO-binding properties of Eme1, with a predominant contribution made by SIM1.

### Eme1 SIMs are required in absence of Rqh1^BLM^

To assess the functional relevance of SIM1 and SIM2, we generated mutant strains harboring the point mutations described in **Figure 5A** in SIM1 (*eme1^SIM1*^*), SIM2 (*eme1^SIM2*^*) or both SIMs (*eme1^SIM1*+SIM2*^*) of Eme1. None of the three *eme1^SIM1*^, eme1^SIM2*^* and *eme1^SIM1*+SIM2*^* mutant strains presented any obvious growth defect or reduced fitness compared to a WT strain in absence of exogenous DNA damage (**Fig 5B and C**) and following CPT treatment (**Fig 5D**). Since *eme1^Δ117^ rqh1Δ* double mutants are non-viable (**Fig 1A**), we next assessed the importance of SIM1 and/or SIM2 for cell viability in absence of Rqh1^BLM^. As shown in **Figures 5E** and **F**, while mutating SIM1 in the *rqh1Δ* background does not reduce colony formation capacities of the resulting *eme1^SIM1*^ rqh1Δ* double mutant compared to the *rqh1Δ* single mutant, it leads to a marked increase in the proportion of elongated and sick cells (**Fig S4A**). In contrast, mutating SIM2 strongly impairs the ability of *eme1^SIM2*^ rqh1Δ* to form viable colonies (**Fig 5E and F**) in addition to causing a strong increase in the number of sick cells (**Fig S4A**). Simultaneously mutating both SIMs did not further impair colony formation capacities compared to *eme1^SIM2*^ rqh1Δ* cells (**Fig 5E and F**). However, it had an additive effect regarding the proportion of elongated and sick cells (**Fig S4A**).

We further looked at cell fitness following chronic exposure to CPT. Loss of Rqh1^BLM^ in *eme1^SIM1*^* mutant slightly exacerbated CPT sensitivity compared to *rqh1Δ*. This effect was slightly more pronounced for *eme1^SIM2*^ rqh1Δ* mutants while the *eme1^SIM1*+SIM2*^ rqh1Δ* mutants displayed the steepest increase in CPT sensitivity compared to *rqh1Δ* (**Fig S4B**). Overall, these data demonstrate that both SIMs contribute to the essential role of the Eme1^1-117^ N-terminal domain in absence of Rqh1^BLM^, with a more prominent contribution made by SIM2.

### SIM2 contributes to DNA damage-induced Eme1 phosphorylation

The importance of the Eme1 SIMs in absence of Rqh1^BLM^ prompted us to assess possible functional ties between the SUMO-binding properties of Eme1 and its phosphorylation. Interestingly, whereas mutating the SIMs had no obvious effect on the phosphorylation profile of Eme1^SIM1*+SIM2*^ throughout the cell cycle (**Fig 6A**), mutating SIM2 alone was sufficient to substantially reduce phosphorylation levels in response to CPT (**Fig 6B**). In contrast, mutating SIM1 had barely any impact. We next investigated which of the fission yeast SUMO E3 ligases, Pli1 and Nse2 (Watts et al., 2007), are required for DNA damage-induced phosphorylation of Eme1. Unexpectedly, we could detect clear DNA damage-induced phosphorylation of Eme1 in both *pli1Δ* and *nse2-SA* CPT-treated cells, as well as in a double mutant *pli1Δ nse2-SA* that lacks both SUMO-E3 ligases activities (**Fig 6C**). These results were replicated in *pmt3Δ* cells that lack SUMO (**Fig 6C**). Thus, the DNA damage-mediated phosphorylation of Eme1 that relies on an intact SIM2 motif does not require a SIM2-SUMO interaction.

**Figure 6:**
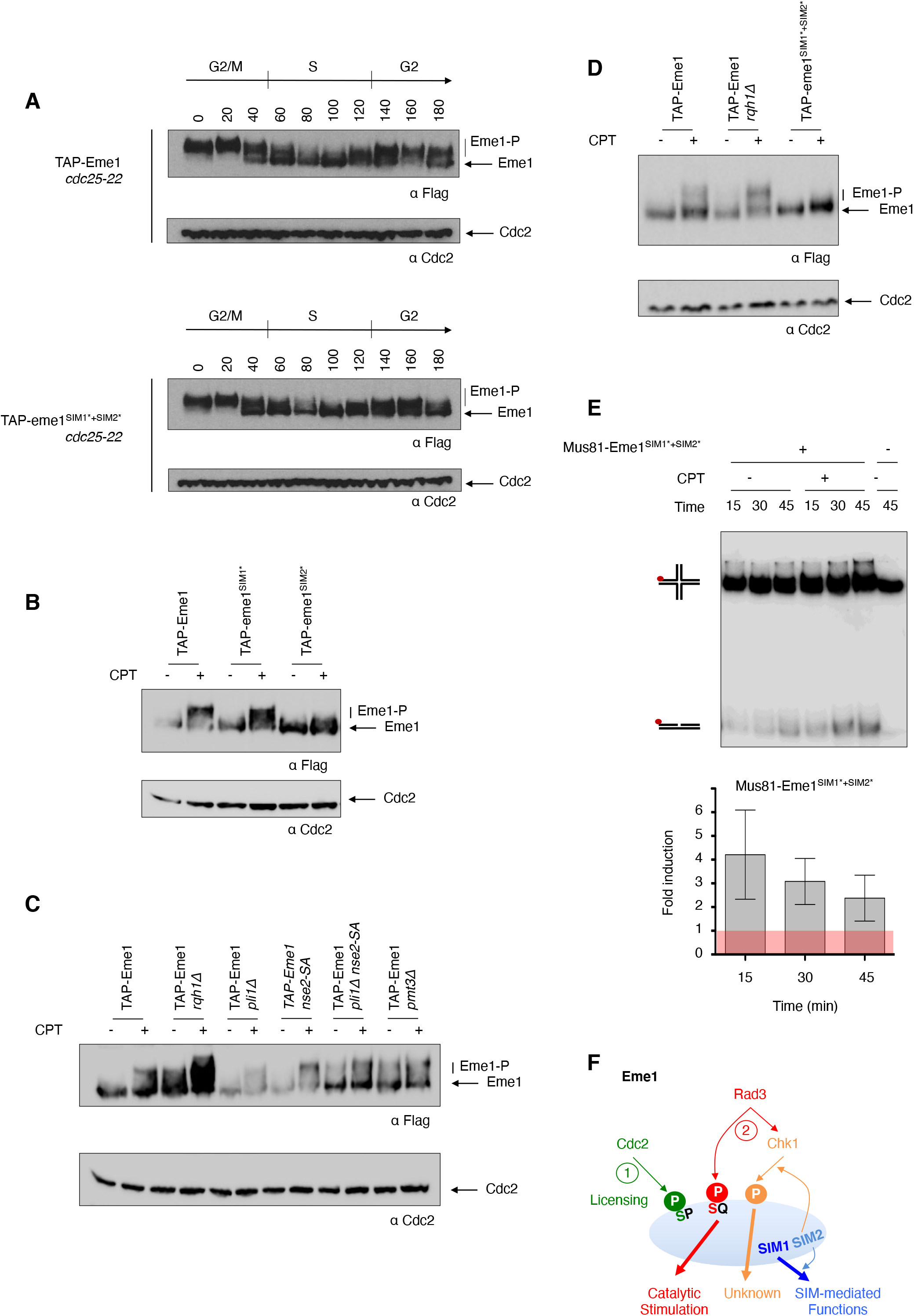
Mutations within the SIMs differently affect Eme1 phosphorylation. A- Cultures from *cdc25-22 TAP-eme1* and *cdc25-22 TAP-eme1^SIM1*+SIM2*^* were synchronized at the G2/M transition and released for one cell cycle. Total proteins were extracted at each indicated time point of the time course and analyzed by Western blot using an antibody raised against the Flag tag of Eme1. Cdc2 is used as a loading control. B- Same as A- for asynchronous cultures from untreated or 40 μM CPT-treated cells of the indicated genotypes. C- Same as A- for asynchronous cultures from untreated or 40 μM CPT-treated cells of the indicated genotypes. D- Same as A- for asynchronous cultures from untreated or 40 μM CPT-treated cells of the indicated genotypes. E- Upper panel: Neutral PAGE showing ^32^P-labeled (red dot) HJs incubated for the indicated times with Mus81-Eme1 complexes recovered from untreated or 40 μM CPT-treated *TAP-eme1^SIM1*+SIM2*^* cells as described in Materials and Methods. Identical amounts of TEV eluates were used in each reaction after normalization of their relative concentration (see Materials and Methods). Lower panel: Average (± s.d.) fold stimulation of HJ resolution by Mus81-Eme1 following CPT-treatment (n=3 independent experiments, see S5). The histogram shows the ratio of the HJ-resolvase activity of Mus81-Eme1 from CPT-treated *TAP-eme1^SIM1*+SIM2*^* cells over that of Mus81-Eme1 from untreated *TAP-eme1^SIM1*+SIM2*^* cells (fold induction). F- Model for the Cdc2-, Rad3^ATR^- and Rad3^ATR^-Chk1-dependent phosphorylation of Eme1 and its SUMO binding properties. The interplay between Eme1 phosphorylation and SUMO-binding properties is detailed in text. Note: TAP- = 2xProtA-TEVsite-2xFlag-

### Eme1 SIMs are not required for Mus81-Eme1 catalytic stimulation

To help decipher whether SIM2-dependent functions are important for phosphorylation of Eme1 by Chk1 or Rad3^ATR^, we investigated whether they contribute to the catalytic stimulation of Mus81-Eme1, which we have shown strickly relies on Rad3^ATR^ and not Chk1 (**Fig 4B,D** and **S3B,D**). Strikingly, the Mus81-Eme1^SIM1*+SIM2*^ complex was efficiently stimulated following CPT-treatment (**Fig 6E and S5**) despite reduced levels of CPT-induced phosphorylation (**Fig 6D**). This demonstrates that the SIM-dependent functions of Eme1 are totally dispensable for direct phosphorylation of Eme1 by Rad3^ATR^ and catalytic stimulation of Mus81-Eme1 and suggests that they are instead key for phosphorylation of Eme1 by Chk1 (**Fig 6F**). Based on these findings and the severe growth defect of an *eme1^SIM2*^ rqh1Δ* double mutant, we propose that phosphorylation of Eme1 by Chk1 represents an additional phosphorylation-based layer of control of Mus81-Eme1 that is non-catalytic yet critical for cell viability in absence of Rqh1^BLM^.

### SUMO-binding capacities and phosphorylation cooperate for cell viability in absence of Rqh1^BLM^

To further assess whether we have unraveled independent layers of regulation of Mus81-Eme1 that all contribute in their own way to cell survival in absence of Rqh1^BLM^, we undertook genetic analyses by combining the mutations that impair the SIMs (i.e. *eme1^SIM1*+SIM2*^*) with those that abrogate the cell-cycle and DNA damage dependent phosphorylations of Eme1 (i.e. *eme1^4SA^*). The resulting_*eme1^SIM1*+SIM2*+4SA^* mutant strain displays a slightly reduced ability to form viable colonies in absence of exogenous stress (**Fig 7A and B**). Remarkably, while *eme1^SIM1*+SIM2*^ rqh1Δ* (**Fig 5E**) and *eme1^4SA^ rqh1Δ* [12] double mutant strains are sick but viable, we were unable to generate a viable *eme1^SIM1*+SIM2*+4SA^ rqh1Δ* strain (**Fig 7C**). This is reminiscent of the synthetic lethal interaction between *eme1^Δ117^* and *rqh1Δ* and demonstrates that the Eme1^1-117^ N-terminal domain is involved in three independent regulatory processes that each make key contributions to the essential functions fulfilled by Mus81-Eme1 in absence of Rqh1^BLM^.

**Figure 7:**
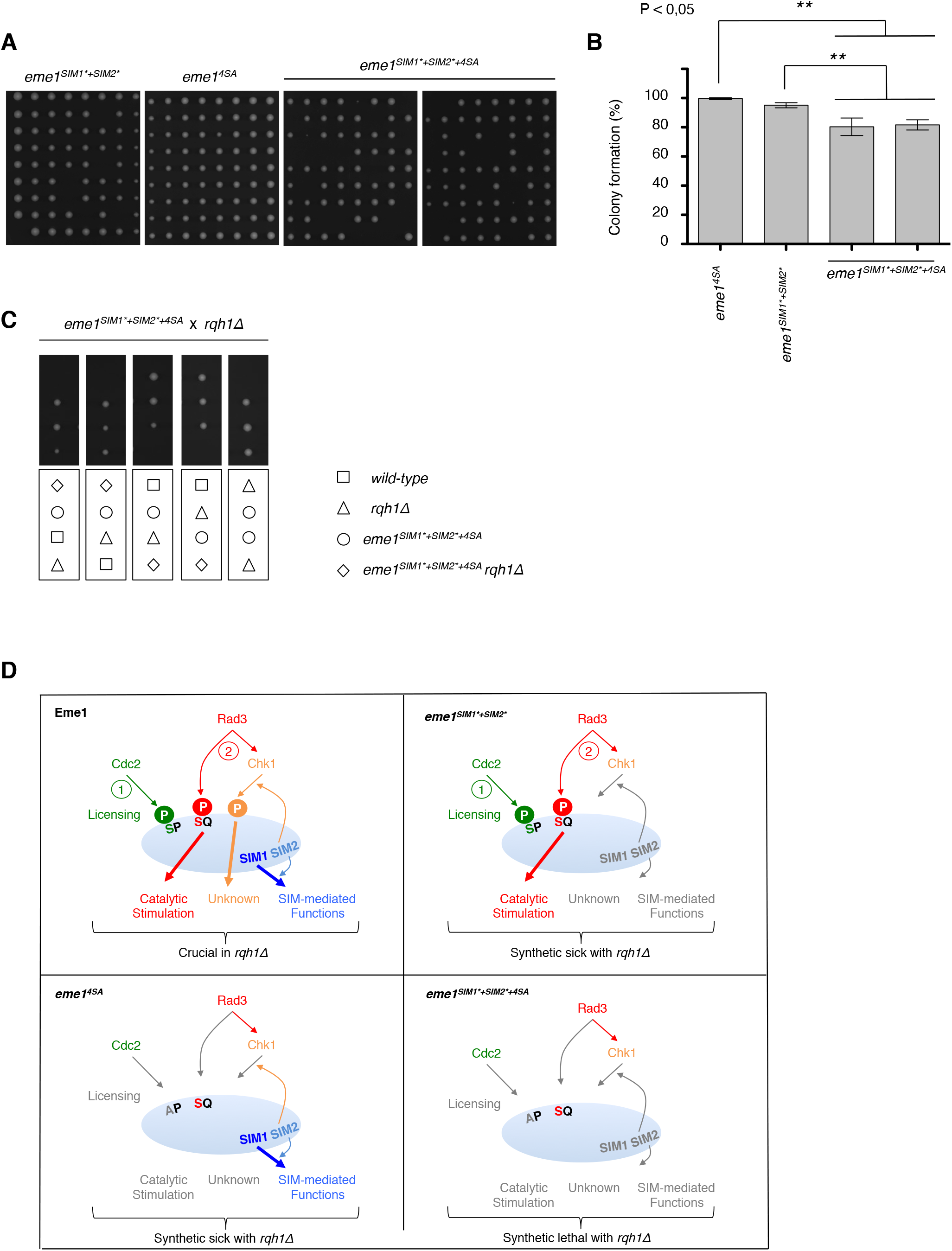
The combined loss of Eme1 phosphorylation and SUMO-binding properties is lethal in absence of Rqh1^BLM^. A- Exponentially growing *eme1^SIM1*+SIM2*^, eme1^4SA^* and *eme1^SIM1*+SIM2*+4SA^* cells were seeded on YES plates by micromanipulation and allowed to grow for 3 days at 32°C. B- Average percentage (± s.d.) of viable colonies (n=3 independent experiments). Statistical significance is measured with one-way ANOVA followed by Tuckey post-test. C- Tetrad analysis of an *eme1^SIM1*+SIM2*+4SA^* x *rqh1Δ* mating, germinated at 30 °C. Boxes below dissections indicate the genotypes of each spore. D- Model depicting the respective contributions of Eme1 phosphorylation and SUMO-binding properties in the phenotype observed in absence of Rqh1^BLM^.

## Discussion

In this study we have identified three independent layers of control of Mus81-Eme1 that are critical in absence of Rqh1^BLM^ (**Fig 7D**). A first layer relies on the phosphorylation of Eme1 by Rad3^ATR^ in response to DNA damage and is critical for the catalytic stimulation of Mus81-Eme1 (**Fig 4E and 7D**). A second layer also relies on the phosphorylation of Eme1 in response to DNA damage, but in this case it is mediated by Chk1 and does not involve the catalytic stimulation of Mus81-Eme1 (**Fig 4E and 7D**). It is noteworthy that both these layers require that Eme1 is first phosphorylated by Cdc2^CDK1^[12]. The third layer relies on newly described SIM1- and SIM2-dependent SUMO-binding properties of Eme1.

In our initial study we had suggested that catalytic stimulation of Mus81-Eme1 in response to DNA damage relied on the phosphorylation of Eme1 by a classical DNA damage checkpoint signaling mechanism and that it was ultimately driven by phosphorylation of Eme1 by Chk1[12]. With the demonstration that it is in fact direct phosphorylation of Eme1 by Rad3^ATR^ and not Chk1 that is critical in that process, we are providing important new insight into a control mechanism of Mus81-Eme1 that turns out to be even more elaborate than initially anticipated. Indeed, in addition to this Rad3-mediated catalytic control, our genetic analyses of the *eme1^SIM2*^* mutant suggest that Chk1-dependent phosphorylation of Eme1, which relies on SIM2 (**Fig 6**), has to be kept in the picture as another functionally important modification for cell fitness in absence of Rqh1^BLM^ (**Fig 5E, S4**). It is noteworthy that all eight Rad3^ATR^ SQ/TQ phosphorylation sites are located before the coiled-coil domain of Eme1 and sit therefore within an intrinsically disordered part of Eme1. This is particularly relevant for the first two phosphorylation sites found in cluster 1 which are the only two within the Eme1^1-117^ domain. They are the ones that also contribute the most to the phosphorylation of Eme1 by Rad3^ATR^ *in vitro*. It will be important to further investigate what structural impact phosphorylation of the N-terminus of Eme1 may have or whether it drives conformational changes by promoting the association with a coactivator and how this stimulates Mus81-Eme1. Hints that the poorly structured N-terminal domain of Eme1 may negatively impact the catalytic activity of Mus81-Eme1 may be seen in the increased *in vitro* activity that results from clipping off large N-terminal domains of human MUS81 and EME1 that are dispensable for the endonuclease function of the complex[14]. This would also be reminiscent of the auto-inhibition of human MUS81-EME1 by the N-terminal Helix-hairpin-Helix domain within MUS81 that is relieved upon association of the N-terminus of MUS81 with SLX4[15]. Cryo-EM studies of XPF-ERCC1 also revealed how conformational changes imposed by DNA binding relieved auto-inhibition by the N-terminal helicase domain of XPF[16]. Members of the XPF-family of SSEs, which all carry their catalytic-relevant functions in the C-terminal part of their subunits, appear to have evolved in a way that provides their N-terminal domains with regulatory functions mediated through controlled conformational changes. Based on our findings, such conformational changes driven by phosphorylation seems like a plausible explanation in the case of Eme1.

By demonstrating that stimulation of Mus81-Eme1 relies on direct phosphorylation of Eme1 by Rad3^ATR^ and not Chk1, we uncouple the catalytic control of Mus81-Eme1 in response to DNA damage from the canonical DNA damage checkpoint signaling pathway. Indeed, we formally ruled out any contribution of Chk1 to the catalytic stimulation of Mus81-Eme1 given that a Mus81-Eme1^8AQ^ mutant complex still undergoes Chk1-mediated phosphorylation of Eme1^8AQ^ but shows absolutely no increased activity in response to DNA damage (**Fig 2E and 4C**). Importantly, phosphorylation of Eme1 by Rad3^ATR^ only occurs when Eme1 has first been phosphorylated by Cdc2^CDK1^, which precludes phosphorylation of Eme1 by Rad3^ATR^ when the DNA replication checkpoint is activated[12]. It will be interesting to further investigate the timing of Mus81-Eme1 stimulation by Rad3^ATR^ in response to DNA damage. Is Mus81-Eme1 stimulated by Rad3^ATR^ before activation of the downstream Chk1 kinase? If so, Chk1-mediated phosphorylation of Eme1 that would occur after stimulation of Mus81-Eme1 could constitute a way to “turn off” Mus81-Eme1 once it has resolved recombination intermediates, maybe by driving its dissociation from chromatin as observed following phosphorylation of Mus81 by Cds1 in response to replication stress[17]. Or, are we instead in a scenario closer to what has been proposed by Smolka and colleagues where Mec1 signaling is uncoupled in *S. cerevisiae* from downstream Rad53 activation to allow direct control of repair enzymes by Mec1 without the need for prolonged cell-cycle arrest[18]?

Another pivotal finding of this study is the identification of two SIMs (SIM1 and SIM2) in the Eme1^1-117^ N-terminal domain that jointly contribute to the newly reported SUMO-binding properties of Eme1, with a predominant contribution of the stronger SUMO-binder SIM1 (**Fig 5A**). It is noteworthy that while mutating the SIMs of Eme1 severely impacts cell viability in absence of Rqh1^BLM^ (**Fig 5E**) it did not lead to any increased sensitivity to CPT (**Fig 5D**). This contrasts with the acute hypersensitivity of *mus81Δ* and *eme1Δ* null cells and indicates that the Eme1 SIM-dependent functions are intimately linked to the control of Mus81-Eme1 in relation to Rqh1^BLM^ functions. This is reminiscent of what we have observed for the *eme1^4SA^* and *eme1^8AQ^* phosphorylation mutants that are unable to undergo Cdc2^CDK1^ or Rad3^ATR^-mediated phosphorylation, respectively([12]and this study). However, we established that the SIM-dependent functions are not related to the catalytic control of Mus81-Eme1 in response to DNA damage. A likely explanation is that instead they contribute to the efficient recruitment and stabilization of Mus81-Eme1 at sites where it is needed to process secondary DNA structures that accumulate in absence of Rqh1^BLM^. In line with this, it was recently proposed that the human SLX4 nuclease scaffold that targets the XPF-ERCC1, MUS81-EME1 and SLX1 SSEs to specific genomic loci contains several SIMs that are involved in its own recruitment to telomeres, PML bodies and DNA damage[19,20]. An alternative and radically opposite explanation could be that the SIMs of Eme1 are involved in a process that negatively controls Mus81-Eme1 by sequestering the nuclease in subnuclear compartments, away from DNA secondary structures such as replication intermediates that could otherwise get opportunistically processed. In absence of Rqh1^BLM^ such structures would accumulate and their premature endonucleolytic processing would be deleterious to the cell. Such compartmentalisation has been observed in human cells with the nucleolar accumulation of MUS81-EME1 in S-phase and its relocalisation out of the nucleolus to sites of DNA damage in replicating cells following UV-irradiation[21].

Having confirmed by yeast two hybrid that SIM1 and SIM2 are *bone fide* SIMs that bind SUMO (**Fig 5A**), the most straightforward assumption would be that they are involved in the control of Mus81-Eme1 by jointly driving the interaction of Eme1 with one or several SUMOylated proteins. They could also drive transient conformational changes through internal interactions between the SIMs of Eme1 and SUMOylated Eme1 or Mus81, which both have been reported to undergo SUMOylation in *S. cerevisiae* and human cells [11,22,23]. Remarkably, we find that SIM1 and SIM2 fulfil independent functions. First of all, mutating the strong SUMO binding SIM1 causes the accumulation of sick cells in absence of Rqh1^BLM^ but it barely impacts colony formation, whereas mutating the much weaker SUMO-binder SIM2 significantly reduces colony formation in addition to causing the accumulation of sick cells. Mutating both SIMs synergistically increased the proportion of sick cells (**Fig 5E, F and S4A**). Furthermore, we also found that SIM2 but not SIM1 is required for the Chk1-dependent phosphorylation of Eme1 in response to DNA damage (**Fig 6B**). These results not only suggest that each SIM fulfils different functions, they also suggest that some of those fulfilled by SIM2 might extend beyond SUMO-binding when putting into perspective the SIM2-specific phenotypes and its relatively poor affinity for SUMO compared to SIM1. In line with this, we found that Eme1 still undergoes DNA damage-triggered phosphorylation in mutant cells that do not produce SUMO (**Fig 6C**). This indicates that the processes that lead to the phosphorylation of Eme1 in response to DNA damage do not involve SUMO, including those that rely on SIM2. It also begs the question of what kind of protein-protein interaction might involve SIM2. While we cannot exclude that the mutations introduced in SIM2 induce structural changes that impact more than just SUMO-binding, there remains the exciting possibility that SIM2 drives the association of Eme1 with a partner that contains a SUMO-like domain (SLD)[24]. The only SLD-containing protein described so far in *S. pombe* is the Rad60 genome stability factor that contains two SLDs, each of which interact with different players of the SUMO pathway[25]. Interestingly, the presumed *S. cerevisiae* Rad60 ortholog Esc2 has been reported to interact with Mus81 via its SLDs and to stimulate the Mus81-Mms4 complex[26]. In addition, Esc2 was recently found to promote the degradation of phosphorylated Mms4[11]. We can exclude similar scenarios involving Rad60 as catalytic stimulation of Mus81-Eme1 still occurs when both SIMs are mutated (**Fig 6E**) and none of the SIMs were found to modulate the levels of Eme1 or phosphorylated Eme1. However, based on such functional promiscuity between Mus81-Mms4 and the SLD-containing Esc2 protein it is tempting to see Rad60 as the ideal candidate for a SIM2-mediated partner of Eme1 that would promote Chk1-dependent phosphorylation of Eme1 in response to DNA damage. This would not be the first example of regulatory processes that involve similar players but different outcomes for Mus81-Eme1 and Mus81-Mms4.

Overall, our findings show that the poorly structured N-terminal domain of Eme1 harbors essential regulatory functions of Mus81-Eme1, the control of which appears to be remarkably more elaborate than initially described. With the demonstration that it relies on three independent regulatory layers that together contribute to the vital functions it fulfils in cells lacking Rqh1^BLM^, we are setting the basis for new lines of investigation that should contribute to a better understanding of the contributions made Mus81-Eme1 in the maintenance of genome stability.

## Materials and Methods

### Fission yeast strains, media, techniques and plasmids

Fission yeast strain genotypes are listed in Supplementary **table 1**. Media and methods for studying *S. pombe* were as described elsewhere[27].

The *eme1* mutants (*eme1^8AQ^, eme1^SIM1*^, eme1^SIM2*^, eme1^SIM1*+SIM2*^, eme1^SIM1*+SIM2*+4SA^*) were generated as follows. The *Eme1* genomic locus from strain PH81 (*h*+, *leu1-32 ura4-D18 TAP-eme1 mus81:13Myc-KanMX6*), which produces N-terminally TAP-tagged Eme1, was subcloned into a TopoTA vector (Invitrogen). Point mutations were introduced on that TopoTA-*EME1* vector by using a Multiprime site-directed mutagenesis kit (Stratagene). Mutations were confirmed by DNA sequencing. The mutated *Eme1* genomic locus from the TopoTA-*Eme1* vector was used as template for PCR using primers forward 5’ - acccatctctcacctaacc - 3’ and reverse 5’-cagtattagcttacagcc - 3’. The PCR fragment was then used to transform strain PH41, in which a URA4 cassette replaces the start codon of EME1 gene. 5’-FOA- resistant clones were selected and confirmed as *TAP-eme1* mutant producing strains by genomic DNA sequencing.

### Cell synchronization

For synchronization of cells by *cdc25-22* block and release, cells containing the temperature-sensitive *cdc25-22* allele were grown to exponential phase at permissive temperature (25 °C) and shifted at restrictive temperature (36 °C) for 3.5 h to arrest the cell cycle in G2. Upon release to permissive temperature (25 °C), the cells synchronously enter the cell cycle. Cells were collected and processed every 20 min. Progression into S phase was monitored microscopically by counting cells that contained septa using calcofluor (Sigma) staining, the appearance of which correlates with S phase.

For DNA damage studies, Bleomycin (Merck) was added to cells arrested at the G2/M transition and further incubated at 36°C for 1,5h.

### Colony formation assay

Fresh cultures were re-seeded on YES plates by micromanipulation and allowed to grow for 3 days at 32°C. At least 3 independent experiments were performed and averaged. Statistical significance (1 way ANOVA and Tuckey test) is displayed on each graph.

### Yeast two-hybrid

Yeast strains were derived from EGY48 (*MATα, ura3, his3, trpl*, and *LexA_op(x6)_* - *LEU2*) containing the pSH18-34 (*LexA_op(x8)_-LacZ, URA3*, and *amp^r^*) plasmid. The fission yeast complementary DNA (cDNA) coding for the first 130 amino acid was cloned into the pJG4-5 (*B42-AD, TRP1*, and *amp^r^*) and the pEG202 [*LexA_(1-202)_DNA-BD, HIS3*, and *amp^r^*] and cotransformed into EGY48 + pSH18-34 strain and plated onto -URA-TRP-HIS medium. To monitor protein interaction, clones were spotted onto 3% Gal-URA-TRP-HIS-LEU and 3% Gal-URA-TRP-HIS-Xgal (80 μg/ml) plates. Plates were incubated at 30°C for 2 to 4 days.

### Protein extraction, Immunoprecipitation and immunoblotting

Cellular lysates were prepared from exponentially growing cell cultures treated with 40μM camptothecin (Sigma) or 5μg/ml bleomycin (Merck). Denatured cell lysates were prepared by TCA precipitation. Cells were suspended in 20% TCA and lysed mechanically using glass beads (Sigma). Following centrifugation, the TCA precipitate was suspended in SDS-PAGE loading buffer (Invitrogen) containing Tris-base. Protein extracts were directly resolved on Tris-acetate 3-8% polyacrylamide NuPAGE gels (Invitrogen). Proteins were transferred to a nitrocellulose Hybond-C membrane (Invitrogen). The membrane was blocked in PBS-T milk 5% and probed by using anti-Flag (Sigma F1804) antibody (1:5,000 dilution), anti Cdc2 (Santa-Cruz sc-53) antibody (1:1000 dilution), anti-tubulin alpha T5168 (Sigma).

### Recombinant MBP-Eme1-Mus81-6-His production and purification

The cDNA of *eme1* and *mus81* were subcloned into the pMBP-parallel1 and pCDFDuet-1 plasmids respectively using in-fusion (Takara) cloning system. A 6-His tag was inserted in frame with mus81 ORF for C-terminal of the protein. Plasmids were co-transformed in Rosetta™ (DE3)pLysS cells. The expression of MBP-Eme1 and Mus81-6His was carried by growing the cells into auto-induced media (Formedium) at 37°C. Cells were harvest at 4°C, resuspended in PBS 1X and kept at −20°C. Lysis buffer 2X (100mM Tris-HCl, pH=8.0, 300mM NaCl, 20% glycerol, 0,2% NP-40, 2mM PMSF, 2mM ß-Mercaptoethanol, protease inhibitor cocktail complete EDTA-free (Roche), 10 mg/mL lysozyme, 20mM imidazole) was added to lysed cells before incubation for 20 min at 4°C. The lysate was cleared by centrifugation before incubation on Ni^2+^ agarose beads (Qiagen) for 2h at 4°C. The beads were washed with 5 volumes of lysis buffer 1X and eluted with lysis buffer 1X supplemented with 250mM imidazole. Eluted complexes were incubated with amylose beads (NEB) for 2h at 4°C. The beads were washed 3 times in lysis buffer 1X and twice in kinase buffer (25mM HEPES-KOH, pH=7.5, 50mMKCl, 10mM MgCl_2_, 10mM MnCl2, 2% glycerol, 0,1% NP-40, 50mM NaF, 1mM Na_3_V0_4_, 50mM ß-glycerophosphate, 1mM DTT). Complexes were eluted in kinase buffer supplemented with 10mM maltose. Proteins were aliquoted, snap-frozen in liquid nitrogen and stored at −80°C for long term storage.

### GFP-Rad3^ATR^ production and *in vitro* kinase assay

GFP-Rad3^ATR^ was transiently expressed from the full-length nmt41 promoter from *cds1Δ chk1Δ rad3Δ* cells treated 2h with bleomycin. Cells pellets were disrupted using a Ball Mill (Retsch) in presence of liquid Nitrogen. Resulting powder was resuspended in 2 volumes/weight of lysis buffer (50mM Tris-HCl, pH=8.0, 500mM NaCl, 10% glycerol, 1% NP-40, 50mM NaF, 1mM Na_3_VO_4_, 50mM ß-glycerophosphate (Sigma), 2mM PMSF, 1mM DTT, protease inhibitor cocktail complete EDTA free (Roche)). Lysates were cleared by centrifugation before incubation on GFP-TRAP agarose beads (Chromotek) at 4°C for 1h. Beads were washed three times with lysis buffer and twice with kinase buffer (25 mM HEPES-KOH, pH=7.5, 50mM KCl, 10mM MgCl_2_, 10mM MnCl2, 2% glycerol, 0,1% NP-40, 50mM NaF, 1mM Na_3_VO_4_, 50mM β-glycerophosphate, 1mM DTT). GFP-Rad3^ATR^ was kept attached on beads for the following kinase assays.

GFP-TRAP-bound GFP-Rad3^ATR^ was resuspended in kinase buffer supplemented 100 μM of cold ATP before addition of 10 μCi g^32^P-ATP and substrates. After 30 min at 30°C, reactions were stopped by the addition of 15 μL of 4X SDS sample buffer. Samples were denatured and resolved by SDS-PAGE electrophoresis. Following Coomassie staining, gel was dried and expose with phosphorimager.

### In vitro nuclease assay

TAP-Eme1 (2xProtA-TEVsite-2xFlag-Eme1) was affinity purified and used in nuclease assays on X12 mobile HJ as previously described[28]. Briefly, cell pellets were resuspended in 1 volume to weight lysis buffer (50mM Tris-HCl, pH=8.0, 150 mM NaCl, 10% glycerol, 0,1% NP-40, 50mM NaF, 50mM β-glycerophosphate (Sigma), 2mM PMSF, protease inhibitor cocktail complete EDTA free (Roche)). Cells were subjected to mechanical lysis using a Ball Mill (MM400 Retsch). For this, usually, 4 to 5 ml of the cell suspension were poured into grinding chambers precooled in liquid nitrogen. The frozen cell pellet was disrupted by 2 agitation runs at 30hz. The resulting powder was resuspended in another 1 volume to weight of lysis buffer and centrifuged. Clear supernatant was loaded onto IgG sepharose beads (Cytiva) for 2h at 4°C (20 μl packed beads used for 4 ml of lysate). After extensive washes, proteins were eluted in presence of 60 μl of AcTEV protease for 1h at RT. In order to determine the relative amount of Mus81-Eme1 between different samples, 3 μl of each TEV eluate was treated with phosphatase before SDS-PAGE and Western blot analysis in order to collapse the Eme1 signal into a single band of dephosphorylated Eme1. The relative intensity of the dephosphorylated Eme1 band was quantified for each sample using the ImageLab software. Dilution folds were calculated and used to bring the concentration of each sample down to that of the least concentrated sample. 3 μl of each normalized sample was used in nuclease assays with ^32^P-labelled DNA substrates as previously described[28].

TCA (Sigma T6399)

Glass beads (Sigma G8772)

Calcofluor (Sigma 18909)

S-(+-)-Camptothecin (Sigma C9911)

Bleomycin (Calbiochem 9041-93-4)

Hydroxyurea (Sigma H8627)

Protease inhibitor Cocktail EDTA-free (Roche 11873580001)

Anti Flag (Sigma F1804)

Anti Cdc2 (Santa-Cruz sc-53)

Anti tubulin (Sigma T5168)

GFP-TRAP magnetic-agarose (Chromotek gtma-20)

Ni-NTA agarose beads (Qiagen 70666-3)

Amylose resin (NEB E8021S)

Phosphatase (NEB P0753S)

IgG Sepharose® 6 Fast Flow (Cytiva 17-0969-01)

AcTEV (Invitrogen 12575015)

## Acknowledgements

Many thanks to Paul Russell, Nick Boddy and Jean-Hugues Guervilly for their critical and careful reading of the manuscript and for their encouragements. “Milles merci” to Charly Chahwan for hinting at the SUMO-related functions of Eme1 a long time ago and for his enthusiastic support. We thank our colleagues of the 3R community at CRCM for their support and stimulating discussions. We thank Samuel Granjeaud for help on statistical analyses. We are grateful to Nick Boddy, Katsunori Tanaka and Benoit Arcangioli for sharing the *nse2-SA, pmt3Δ* and *pli1Δ* mutant strains, respectively. This work was supported by grants from Agence Nationale de la Recherche (ANR-10BLAN-1512-01) and Institut National du Cancer (INCa-PLBio2016-159 and INCa-PLBio2019-152) awarded to PHLG and a Dotation Jeune Chercheur of the Institut National de la Santé et de la Recherche Médicale awarded to PMD. CG was a recipient of fellowships awarded by the Fondation pour la Recherche Médicale (FRM grant number ECO20170637468) and the Fondation ARC (grant ARCDOC42020010001276).

## Author contributions

PMD, CG and PHLG conceived the experiments. PMD and CG designed and carried out all genetic analyses. PMD and CG undertook the *in silico* analyses of Eme1 primary sequence. PMD focused primarily on the SUMO-related functions of Eme1 whilst CG worked primarily on the phosphorylation of Eme1 by Rad3^ATR^. CG designed and performed all *in vitro* nuclease assays with help from SS. The *in vitro* kinase assays were designed and set up by CG. SC and SS carried out Y2H assays. SS provided technical assistance for routine molecular biology and cloning techniques. PMD and PHLG wrote the paper and all authors read and corrected the manuscript.

## Supplementary Figure legends

**Figure S1:**
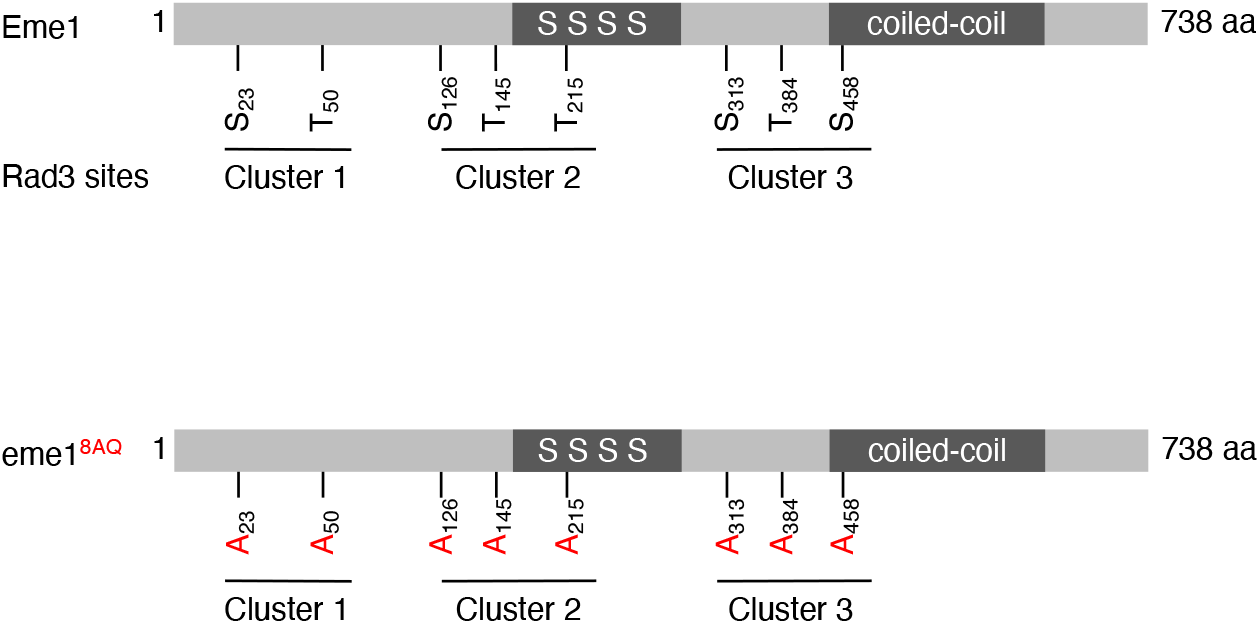
The Rad3^ATR^-consensus sites are widespread throughout Eme1 sequence and were subdivided in three clusters (Upper panel). All Rad3^ATR^-consensus sites are mutated in Alanine to generate eme1^8AQ^ mutant (Lower panel).

**Figure S2:**
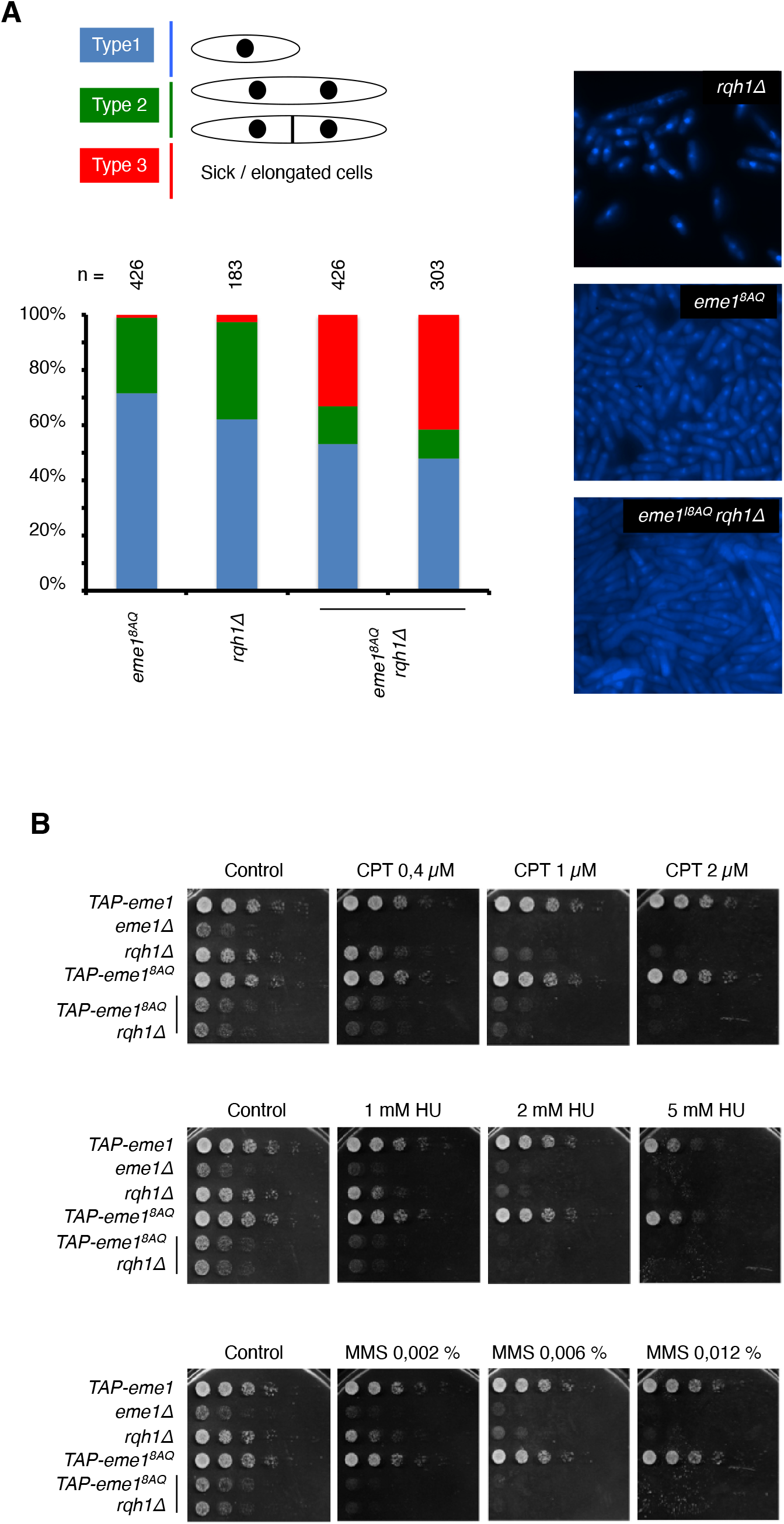
A- Exponentially growing cultures of *eme1^8AQ^, rqh1Δ* and *eme1^8AQ^ rqh1Δ* cells were heated-fixed and observed by fluorescent-microscopy using DAPI staining. Cells were classified based on their morphologies. Small G2 cells (Type 1), elongated binucleated and/or septated cells (Type 2) and sick cells (Type 3). B- Five-fold dilutions of cells with the indicated genotype were plated on medium supplemented or not with the indicated concentrations of CPT, HU and MMS followed by incubation at 30 °C.

**Figure S3:**
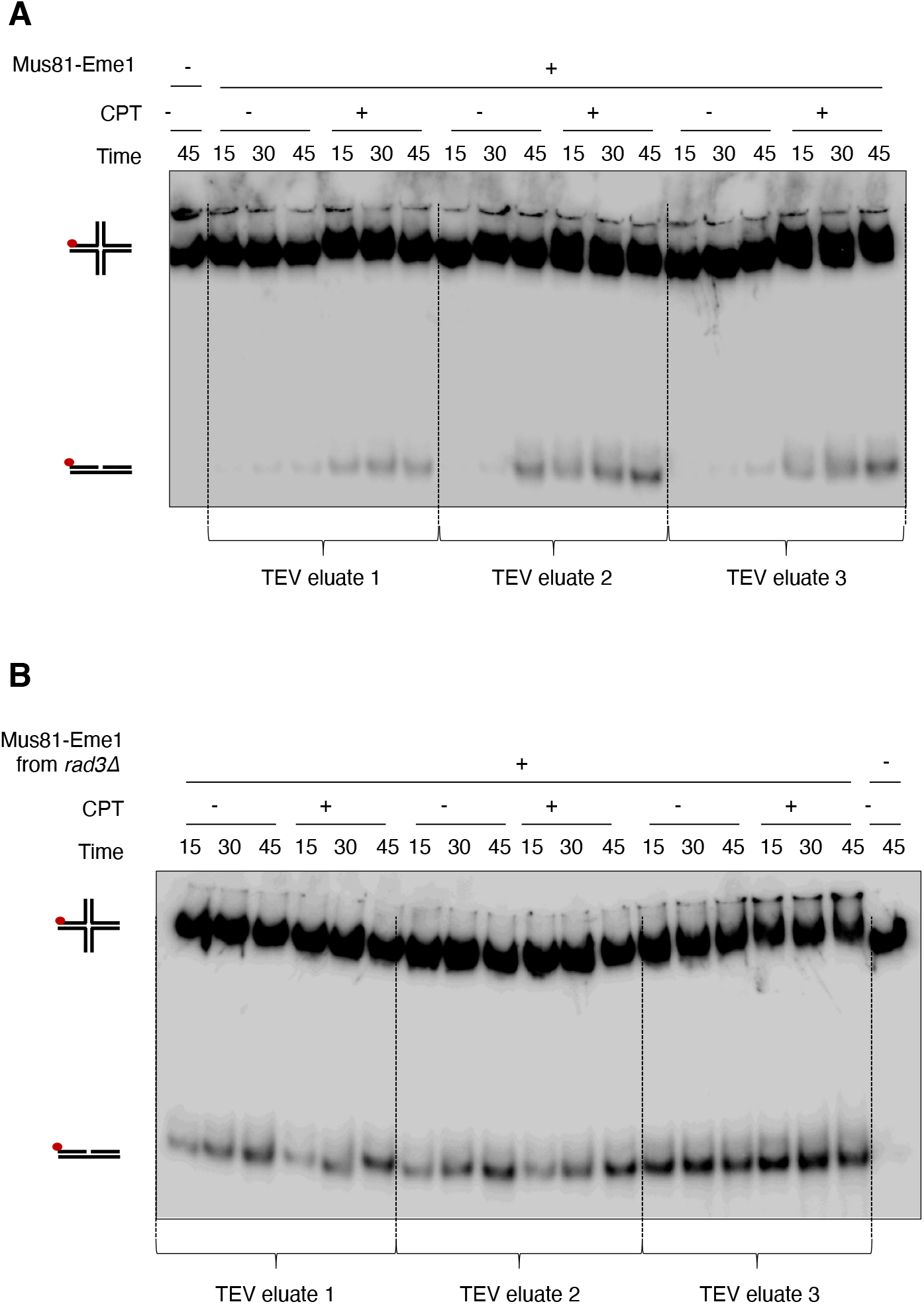
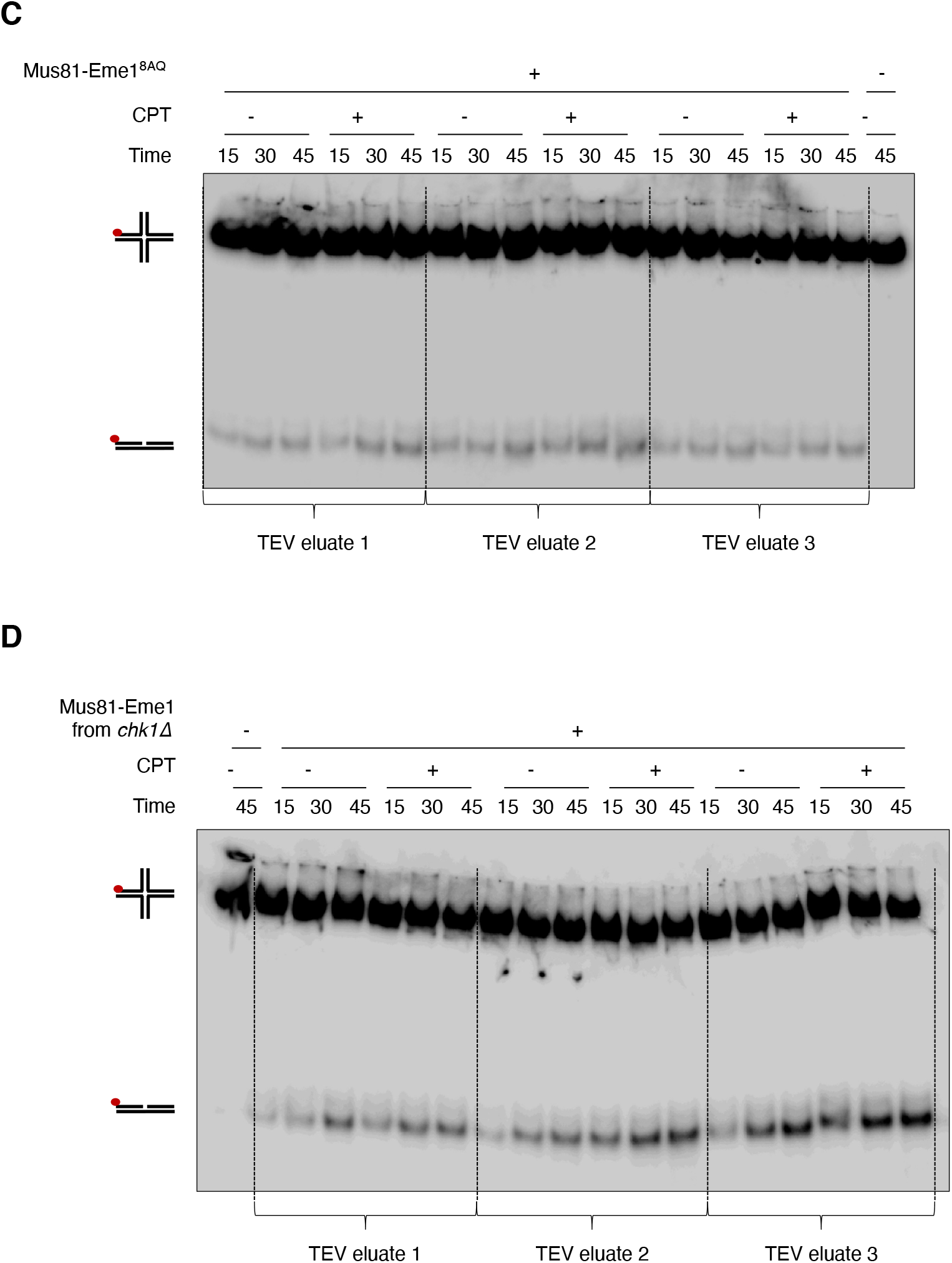
A- Neutral PAGE showing ^32^P-labeled (red dot) HJs incubated for the indicated times with Mus81-Eme1 complexes recovered from untreated or 40 μM CPT-treated *TAP-eme1* (“wild type”) cells as described in Materials and Methods. Identical amounts of TEV eluates were used in each reaction after normalization of their relative concentration (see Materials and Methods). B- Same as A- but with Mus81-Eme1 from untreated or 40 μM CPT-treated *TAP-eme1 rad3Δ* cells. C- Same as A- but with Mus81-Eme1 from untreated or 40 μM CPT-treated *TAP-eme1 eme1^8AQ^* cells. D- Same as A- but with Mus81-Eme1 from untreated or 40 μM CPT-treated *TAP-eme1 chk1Δ* cells. Note: TAP- = 2xProtA-TEVsite-2xFlag-

**Figure S4:**
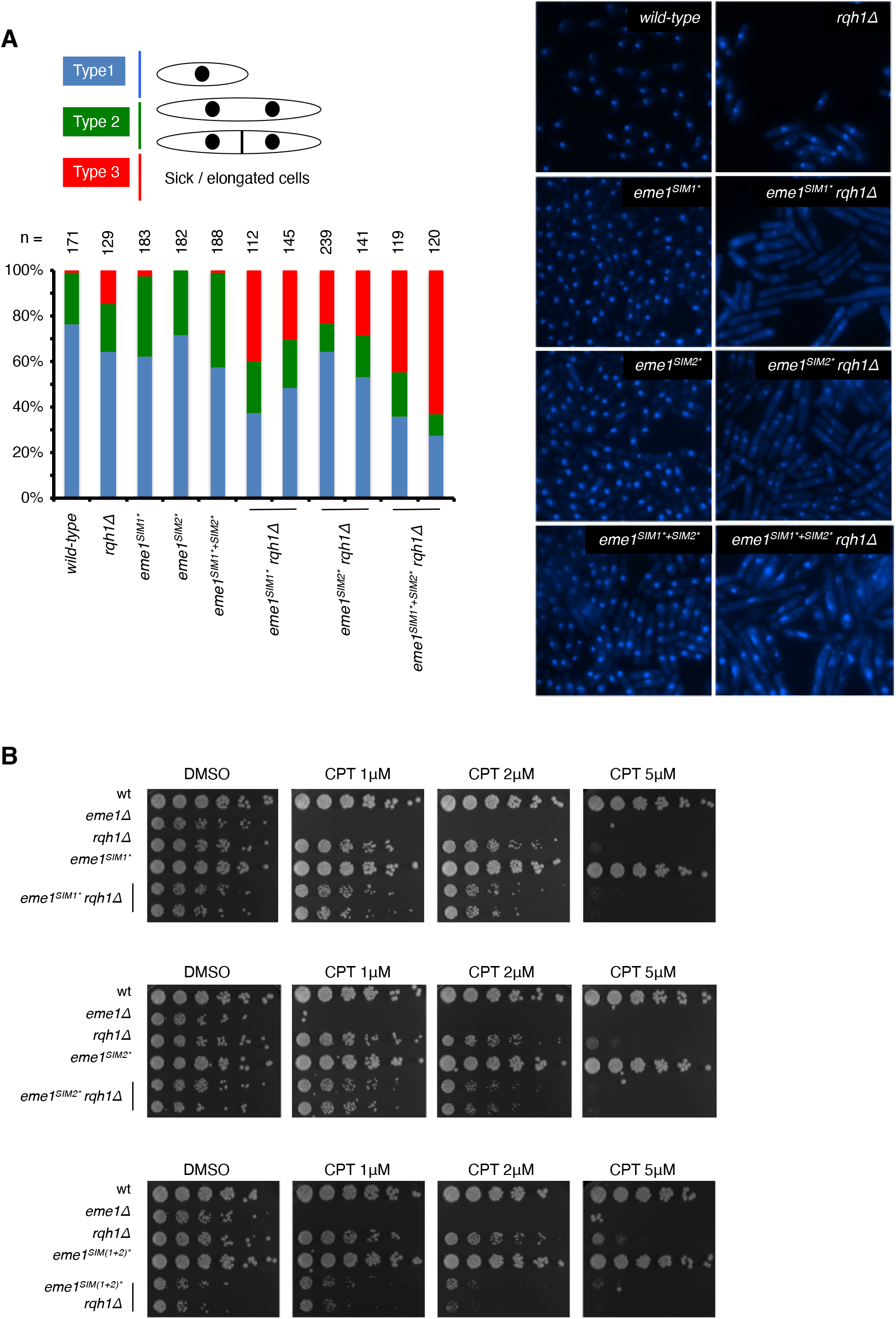
A- Cells from exponentially growing cultures of the indicated genotypes were heated-fixed and observed by fluorescent-microscopy using DAPI staining (same as S2), counted and sorted depending on their morphologies. B- Five-fold dilutions of cells with the indicated genotype were plated on medium supplemented or not with the indicated concentrations of CPT followed by incubation at 30 °C.

**Figure S5:**
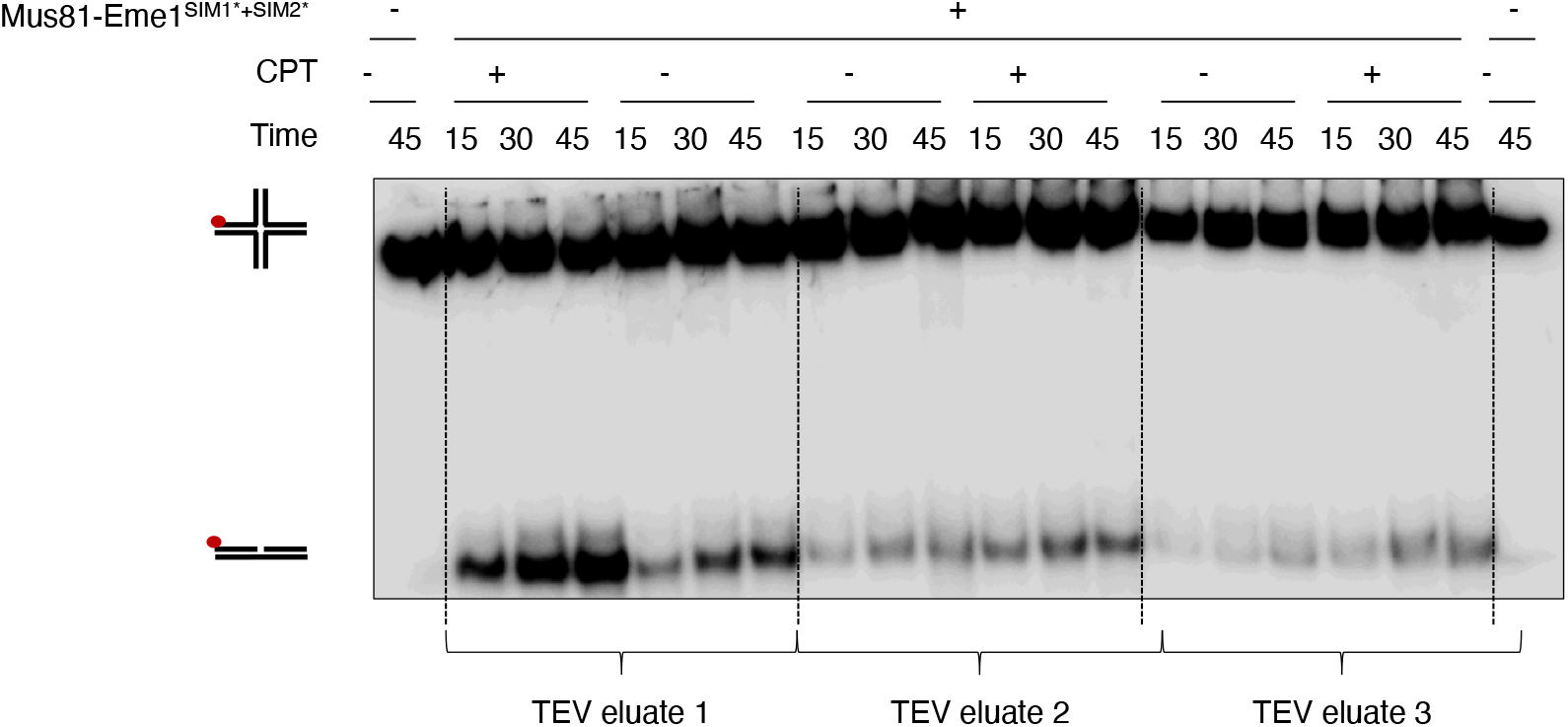
Neutral PAGE showing ^32^P-labeled (red dot) HJs incubated for the indicated times with Mus81-Eme1 complexes recovered from untreated or 40 μM CPT-treated *TAP-eme1^SIM1*+SIM2*^* cells as described in Materials and Methods. Identical amounts of TEV eluates were used in each reaction after normalization of their relative concentration (see Materials and Methods). Note: TAP- = 2xProtA-TEVsite-2xFlag-

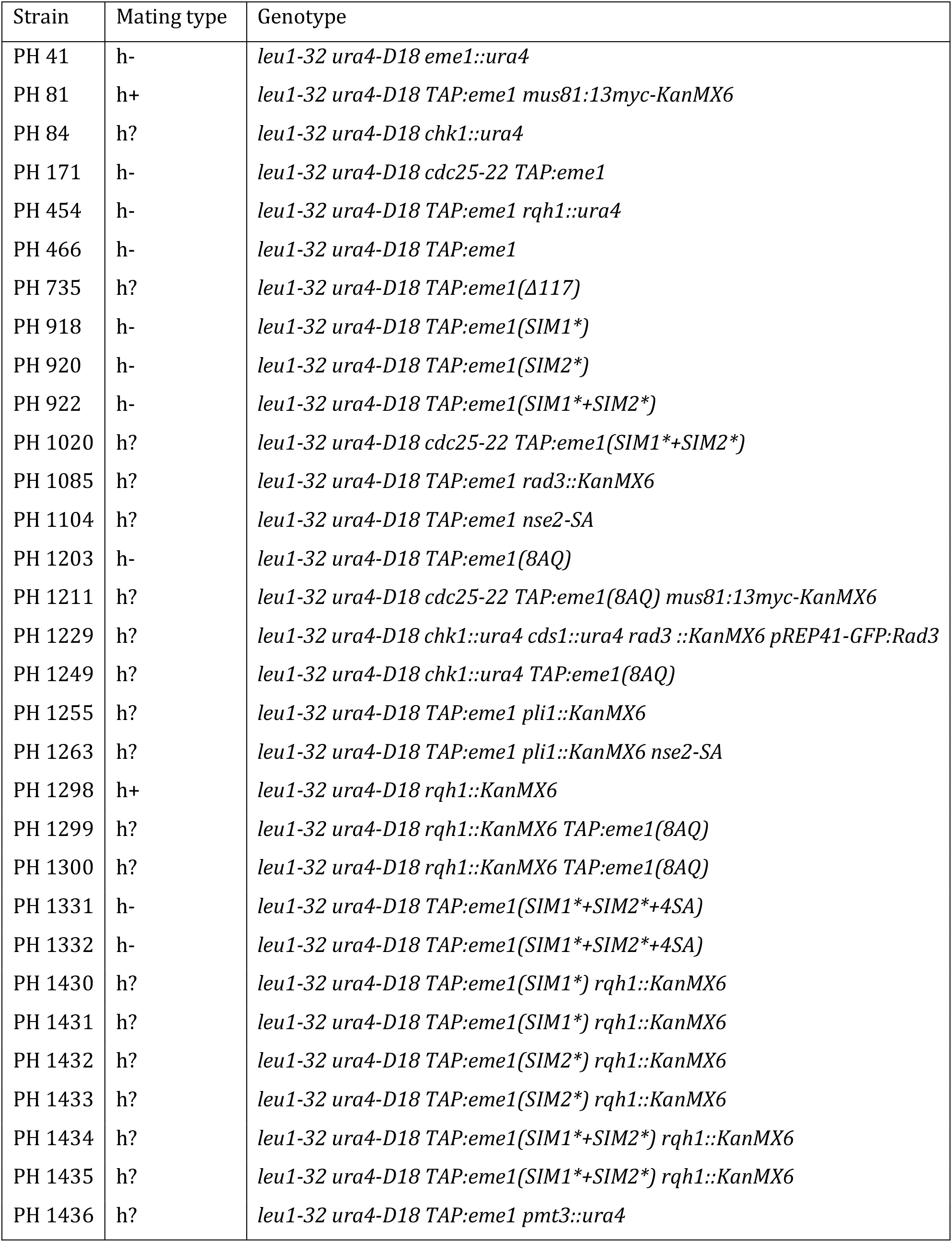

| Strain | Mating type | Genotype |
| --- | --- | --- |
| PH 41 | h- | *leu1-32 ura4-D18 eme1::ura4* |
| PH 81 | h+ | *leu1-32 ura4-D18 TAP:eme1 mus81:13myc-KanMX6* |
| PH 84 | h? | *leu1-32 ura4-D18 chk1::ura4* |
| PH 171 | h- | *leu1-32 ura4-D18 cdc25-22 TAP:eme1* |
| PH 454 | h- | *leu1-32 ura4-D18 TAP:eme1 rqh1::ura4* |
| PH 466 | h- | *leu1-32 ura4-D18 TAP:eme1* |
| PH 735 | h? | *leu1-32 ura4-D18 TAP:eme1(Δ117)* |
| PH 918 | h- | *leu1-32 ura4-D18 TAP:eme1(SIM1\*)* |
| PH 920 | h- | *leu1-32 ura4-D18 TAP:eme1(SIM2\*)* |
| PH 922 | h- | *leu1-32 ura4-D18 TAP:eme1(SIM1\*+SIM2\*)* |
| PH 1020 | h? | *leu1-32 ura4-D18 cdc25-22 TAP:eme1(SIM1\*+SIM2\*)* |
| PH 1085 | h? | *leu1-32 ura4-D18 TAP:eme1 rad3::KanMX6* |
| PH 1104 | h? | *leu1-32 ura4-D18 TAP:eme1 nse2-SA* |
| PH 1203 | h- | *leu1-32 ura4-D18 TAP:eme1(8AQ)* |
| PH 1211 | h? | *leu1-32 ura4-D18 cdc25-22 TAP:eme1(8AQ) mus81:13myc-KanMX6* |
| PH 1229 | h? | *leu1-32 ura4-D18 chk1::ura4 cds1::ura4 rad3::KanMX6 pREP41-GFP:Rad3* |
| PH 1249 | h? | *leu1-32 ura4-D18 chk1::ura4 TAP:eme1(8AQ)* |
| PH 1255 | h? | *leu1-32 ura4-D18 TAP:eme1 pli1::KanMX6* |
| PH 1263 | h? | *leu1-32 ura4-D18 TAP:eme1 pli1::KanMX6 nse2-SA* |
| PH 1298 | h+ | *leu1-32 ura4-D18 rqh1::KanMX6* |
| PH 1299 | h? | *leu1-32 ura4-D18 rqh1::KanMX6 TAP:eme1(8AQ)* |
| PH 1300 | h? | *leu1-32 ura4-D18 rqh1::KanMX6 TAP:eme1(8AQ)* |
| PH 1331 | h- | *leu1-32 ura4-D18 TAP:eme1(SIM1\*+SIM2\*+4SA)* |
| PH 1332 | h- | *leu1-32 ura4-D18 TAP:eme1(SIM1\*+SIM2\*+4SA)* |
| PH 1430 | h? | *leu1-32 ura4-D18 TAP:eme1(SIM1\*) rqh1::KanMX6* |
| PH 1431 | h? | *leu1-32 ura4-D18 TAP:eme1(SIM1\*) rqh1::KanMX6* |
| PH 1432 | h? | *leu1-32 ura4-D18 TAP:eme1(SIM2\*) rqh1::KanMX6* |
| PH 1433 | h? | *leu1-32 ura4-D18 TAP:eme1(SIM2\*) rqh1::KanMX6* |
| PH 1434 | h? | *leu1-32 ura4-D18 TAP:eme1(SIM1\*+SIM2\*) rqh1::KanMX6* |
| PH 1435 | h? | *leu1-32 ura4-D18 TAP:eme1(SIM1\*+SIM2\*) rqh1::KanMX6* |
| PH 1436 | h? | *leu1-32 ura4-D18 TAP:eme1 pmt3::ura4* |

